# Hepatitis C Virus Infects and Perturbs Liver Stem Cells

**DOI:** 10.1101/2021.10.26.465357

**Authors:** Nathan L Meyers, Tal Ashuach, Danielle E Lyons, Camille R Simoneau, Ann L Erickson, Mehdi Bouhaddou, Thong T. Nguyen, Mir M Khalid, Taha Y Taha, Vaishaali Natarajan, Jody L Baron, Norma Neff, Fabio Zanini, Tokameh Mahmoud, Stephen R Quake, Nevan J Krogan, Stewart Cooper, Todd C McDevitt, Nir Yosef, Melanie Ott

**Affiliations:** Gladstone Institute of Virology, San Francisco, CA, USA; Department of Electrical Engineering and Computer Science and Center for Computational Biology, University of California Berkeley, Berkeley, CA, USA; California Pacific Medical Center Research Institute, San Francisco, CA, USA; Cellular and Molecular Pharmacology, University of California, San Francisco, CA, USA; Gladstone Institute of Data Science and Biotechnology, San Francisco, CA, USA; Quantitative Biosciences Institute, University of California, San Francisco, CA, USA; Gladstone Institute of Cardiovascular Disease, San Francisco, CA, USA; Department of Medicine, University of California San Francisco, San Francisco, CA, USA; Chan Zuckerburg Biohub, San Francisco, CA, USA; Department of Bioengineering, Stanford University, Stanford, CA, USA; Department of Biochemistry, Erasmus University Medical Center, Rotterdam, the Netherlands; Bioengineering & Therapeutic Sciences, University of California San Francisco, San Francisco, CA, USA; Ragon Institute of Massachusetts General Hospital, MIT and Harvard, Cambridge, MA, USA; Chan Zuckerberg Biohub Investigator, San Francisco, CA, USA; Liver Center, University of California San Francisco, San Francisco, CA, USA

**Keywords:** hepatitis c virus, organoid, liver disease, stem cell, single-cell RNA-sequencing, chronic infection, hepatocellular carcinoma

## Abstract

Hepatitis C virus (HCV) is the leading cause of death from liver disease. How HCV infection causes lasting liver damage and increases cancer risk beyond viral clearance remains unclear. We identify bipotent liver stem cells as novel targets for HCV infection, and their erroneous differentiation as the potential cause of impaired liver regeneration and cancer development. We show 3D organoids generated from liver stem cells from actively HCV-infected individuals carry replicating virus and maintain low-grade infection over months. Organoids can be infected with a primary HCV isolate. Virus-inclusive single-cell RNA-sequencing uncovered extensive transcriptional reprogramming in HCV^+^ cells supporting hepatocytic differentiation, cancer stem cell development and viral replication while stem cell proliferation and interferon signaling are disrupted. Our data adds a pathogenesis factor – infection of liver stem cells – to the biology of HCV infection that explains persistent liver damage and enhanced cancer risk through an altered stem cell state.

## Introduction

Hepatitis C virus (HCV) is a leading cause of hepatocellular carcinoma (HCC) (Koike, 2014). Approximately 71 million individuals are chronically infected with HCV and at risk of liver disease, including fibrosis, steatosis, cirrhosis, and HCC (Modin et al., 2019). Direct-acting antivirals effectively eradicate HCV (Houghton, 2019, Wang et al., 2019), but they do not reverse terminal liver disease, and treated individuals remain at risk for HCC (Wang et al., 2019, Hsu et al., 2019). In the healthy liver, resident bipotent adult stem cells regenerate the tissue after injury by giving rise to hepatocytes and ductal cells (Nio et al., 2017, Lo and Ng, 2013, Qiu et al., 2018). The persistence of liver disease after virus eradication may therefore indicate that these stem cells have sustained durable damage that prevents them from regenerating the tissue lost to infection. Indeed, in chronically infected HCV patients, chronic intrahepatic inflammation activates the proliferation of liver stem cells (Prakoso et al., 2014), which is linked to the development and progression of fibrosis (Helal et al., 2018) and are at the origin of primary liver cancers (Lo and Ng, 2013). These findings confirm that liver stem cells are altered by infection, in ways that could lead to lasting liver damage in patients with chronic HCV infection even after treatment with direct-acting antivirals. But whether durable alterations of the liver stem cells result solely from inflammation or also from direct infection remains unclear.

Demonstrating that HCV can infect liver stem cells is challenging. HCV and especially primary viral isolates are difficult to culture *ex vivo*. Long-term replication of HCV in adult or fetal primary human liver cells is hampered by the progressive de-differentiation of liver cells in culture and a robust interferon response in primary human liver cells preventing viral replication (Ploss et al., 2010, Andrus et al., 2011, Marukian et al., 2011, Wu et al., 2012, Schobel et al., 2018, Wu et al., 2018). Many studies use cell-culture-adapted strains in a Huh 7-derived hepatoma cell line with defective interferon signaling. However, these cells lack proper cell polarity and display abnormal lipid metabolism (Wakita et al., 2005, Steinmann and Pietschmann, 2013, Catanese and Dorner, 2015, Douam and Ploss, 2015). Hepatocyte-like cells from human induced pluripotent stem cells (iPSCs) would offer a more physiological model (Wang et al., 2019, Wu et al., 2012, Carpentier et al., 2014, Schwartz et al., 2012, Schobel et al., 2018). However, iPSCs are refractory to HCV infection due to high intrinsic expression of interferon-stimulated genes (ISGs). The same problem arises with fetal hepatic progenitor cells (Wu et al., 2012, Schobel et al., 2018, Wu et al., 2018).

To overcome these barriers, we examined adult liver stem cells of individuals infected with HCV. Stem cells with bipotent characteristics analogous to those of hepatic progenitor cells can be cultured *ex vivo* in a 3D organoid form (Broutier et al., 2016, Huch et al., 2015). 3D liver organoids maintain biopotency, with the ability to differentiate into hepatocyte- and cholangiocyte-like cells, cell polarity and reproduce *in vitro* the *in vivo* phenotype of several liver diseases (Broutier et al., 2016, Broutier et al., 2017, Huch et al., 2015). We therefore generated bipotent liver stem cell organoids from HCV-infected individuals. We found that the organoids carried HCV and successfully maintained HCV infection in long-term culture, opening the intriguing possibility that direct infection of liver stem cells contributes to the hepatopathogenesis of HCV infection.

## Results

### Liver organoids from infected and uninfected individuals are phenotypically similar

In order to determine if adult liver stem cell organoids could be derived from HCV+ donors, we generated organoids from liver resections from six donors (Table 1) as described (Broutier et al., 2016, Huch et al., 2015). Three donors had no viral infection (NV) but various forms of metastatic cancers. Three donors were viremic, presented with different HCV genotypes (HCV1–3), and had virus-associated HCC. Donors were 63–70 years old with one female and two males in each group.

**Table 1.**
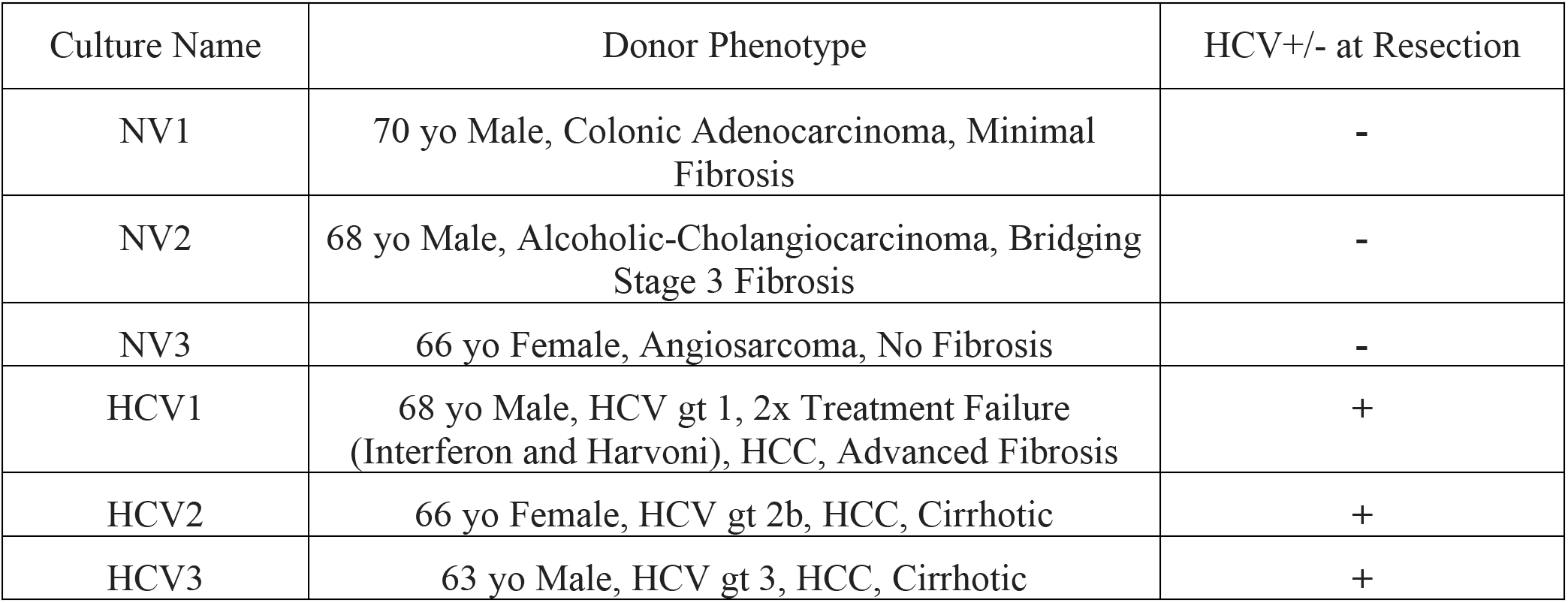
Liver Organoids Grow from Non-Viral and HCV-Infected Individuals. Liver organoids generated from six donors, using single-cell digests from patient liver resections. Three non-viral (NV) patients with various metastatic cancers that necessitated liver resection. Three donors were HCV patients that were viremic at the time of resection. Their HCV genotypes were known and match donor number (i.e., HCV1, 2, 3).

| Culture Name | Donor Phenotype | HCV+/- at Resection |
| --- | --- | --- |
| NV1 | 70 yo Male, Colonic Adenocarcinoma, Minimal Fibrosis | - |
| NV2 | 68 yo Male, Alcoholic-Cholangiocarcinoma, Bridging Stage 3 Fibrosis | - |
| NV3 | 66 yo Female, Angiosarcoma, No Fibrosis | - |
| HCV1 | 68 yo Male, HCV gt 1, 2x Treatment Failure (Interferon and Harvoni), HCC, Advanced Fibrosis | + |
| HCV2 | 66 yo Female, HCV gt 2b, HCC, Cirrhotic | + |
| HCV3 | 63 yo Male, HCV gt 3, HCC, Cirrhotic | + |

NV and HCV^+^ organoids grew in a progenitor state in expansion medium (EM) with similar lifespans of ~4 months in culture (Fig. 1A, S1). They differentiated towards a hepatocyte-like fate when switched to differentiation medium (DM) (i.e., Wnt and Notch agonists were replaced with Notch inhibitor DAPT, dexamethasone, and TGFβ & & bone morphogenic protein (BMP) agonists (Broutier et al., 2017, Huch et al., 2015)). Upon differentiation, proliferation slowed, organoids became opaque and transitioned from an epithelial monolayer to a pseudostratified epithelium (Broutier et al., 2017, Huch et al., 2015). No phenotypic differences were observed between NV and HCV^+^ organoids (Fig. 1A, D). Differentiation of both NV and HCV^+^ organoids showed similar induction of mRNA expression of several hepatocyte markers, Albumin (ALB) and cytochrome P450 enzymes (CYP3A4, and CYP2B6), although to a lesser extent than primary human hepatocytes, and decreased expression of the stem-cell marker LGR5 (Fig. 1B). Transcripts for HCV entry factors including Cluster of Differentiation 81 (CD81), Occludin (OCLDN), Claudin-1 (CLDN1), and Scavenger Receptor Class B Type I (SR-B1) were more highly expressed in organoids than in primary human hepatocytes and at similar levels before and after differentiation (Fig. 1C). CD81 and Occludin were properly expressed and located at tight junctions and the apical membrane (Fig. 1D). Differentiated organoids from both groups expressed albumin and the hepatocyte nuclear factor 4α (HNF4A) proteins, and were polarized with the apical membrane, marked by Zonula occludens-1 (ZO-1), facing the organoid lumen (Fig. 1D). These results demonstrate that liver organoids can be generated from HCV^+^ donors, and they display similar organoid forming properties as those derived from HCV^−^ donors.

**Figure 1.**
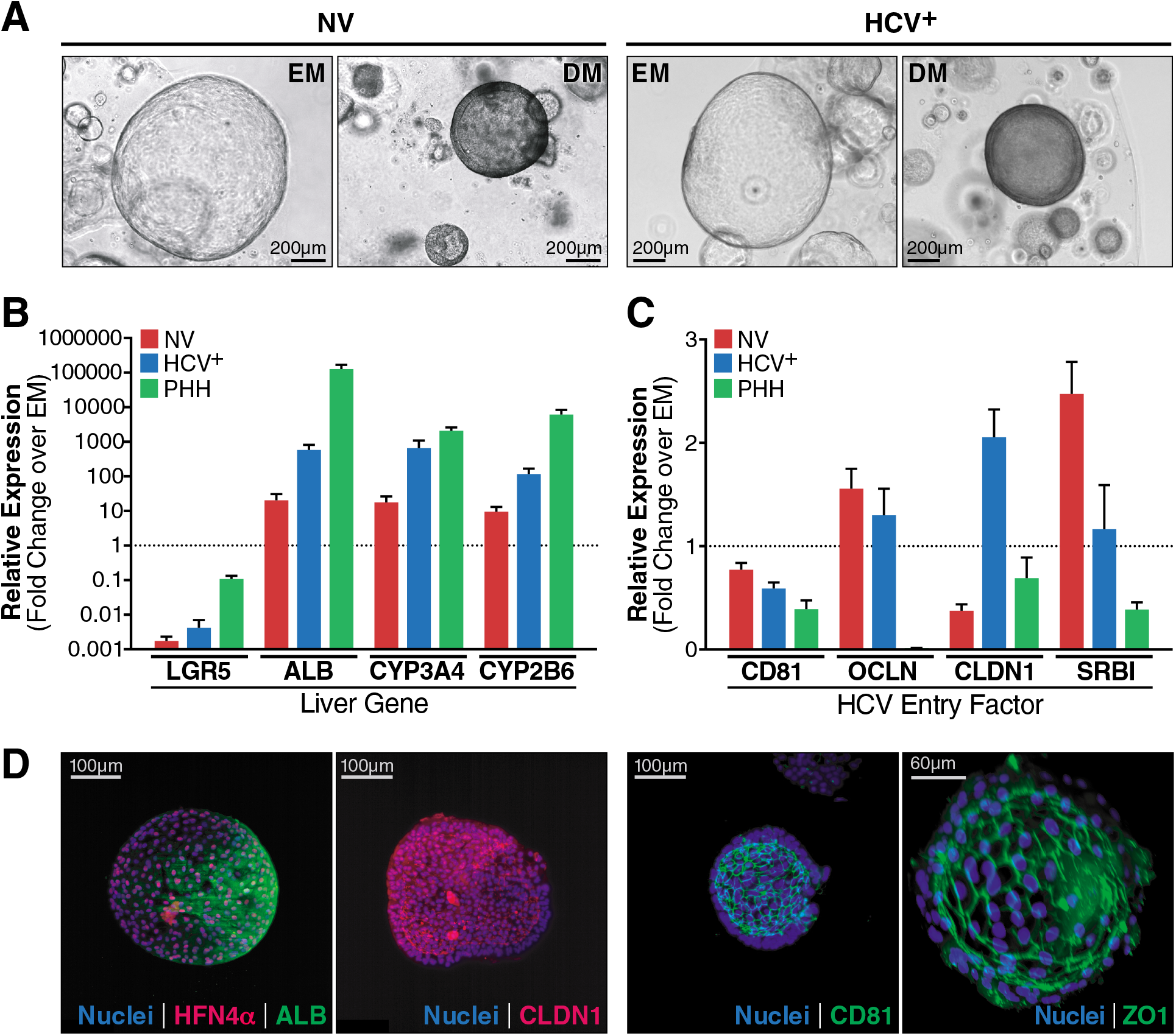
Liver organoids grow from HCV-infected individuals and show similar differentiation potential. **(A)** Representative brightfield microscopy images of liver organoids grown from uninfected (NV) or HCV^+^ donors are shown in the stem-cell (EM) and differentiated (DM) states. Organoids are morphologically distinct in EM vs DM states, but each state was morphologically identical across all six NV and HCV+ donors. **(B)** qPCR quantification of hepatocyte stem-cell marker LGR5, and hepatocyte markers ALB, CYP3A4, and CYP2B6 in DM organoids from three NV donors and three HCV^+^ donors, and in two primary hepatocyte samples relative to EM. For each gene, data was pooled from n≥2 biological replicates per organoid or hepatocyte donor and represented as mean ± SD. Transcript expression was normalized to 18S and plotted as a fold change over the gene’s expression in EM (ΔΔC_T_) which was set to 1 and is marked by a dotted line. Fold change was plotted on a log10 scale. **(C)** qPCR quantification of HCV entry markers, CD81, OCLN, CLDN1, and SR-B1 in the same samples as in (B). EM expression levels were set to one (marked by a dotted line), and fold change in DM was plotted on a linear scale. **(D)** Representative light-sheet microscopy images are shown for differentiated liver organoids (DM) stained for hepatocyte markers HNF4α and ALB, HCV entry factors CLDN1 and CD81, or apical membrane marker ZO1.

### Single-cell transcriptional profiling points to persistent stem cell infection

To identify differences between NV and HCV^+^ organoids, we performed single-cell RNA-sequencing (scRNA-seq) on EM organoids from two NV and three HCV^+^ donors using the 10X Genomics Drop-Seq protocol (Fig. 2, Data S1) (Zheng et al., 2017). We analyzed the combined data sets (Table S1) using scVI, a deep generative model for analyzing scRNA-seq data, which accounts for limited sensitivity and batch effects (Lopez et al., 2018), followed by clustering and annotation using Vision (DeTomaso et al., 2019). We identified 11 clusters containing cells from the five organoid cultures. Cells from different donors were well mixed across clusters, except for cells from the HCV3 donor, which were less well distributed and mostly enriched in clusters 10 and 11 (Fig. 2).

**Figure 2.**
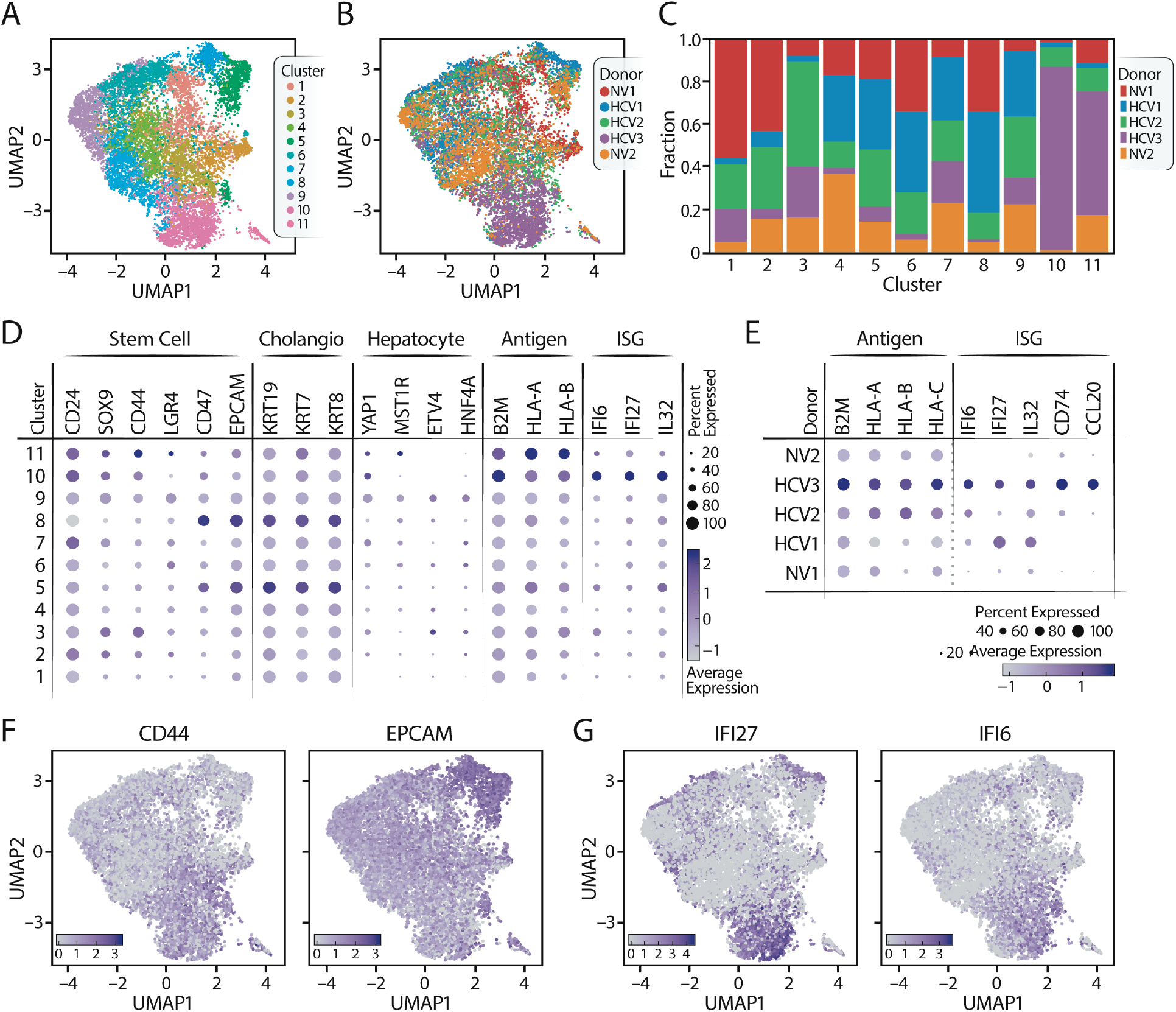
Liver organoids derived from HCV^+^ donors upregulate antigen presentation and interferon-stimulated genes. **(A,B)** UMAPs of 3’ scRNA-seq of EM organoids from five donors are plotted with cells labeled by cluster (A) and donor (B). **(C)** A Stacked bar plot shows the donor composition of each cluster. Clusters are shown on the x-axis and the fraction of cells from each donor within that cluster is plotted on the y-axis. Donor color matches the colors used in (A). **(D)** A dot-plot shows expression (as pseudolog) by cluster of cancer stem-cell markers (stem), ductal, hepatocyte, and antiviral markers, including antigen presentation genes (Antigen) and interferon-stimulated genes (ISGs). **(E)** A dot-plot shows gene expression by donor (as pseudolog) for antigen presentation genes and ISGs. **(F,G)** UMAPs for gene expression (as pseudolog) of two of the stem-cell markers (CD44, EPCAM) (F) and two of the ISGs (IFI27, IFI6) (G). These plots superimpose the UMAP plot in (A) with the relative expression levels of a given gene.

Since EM organoids maintain a stem-cell-like state, we focused on expression of stem-cell markers (Fig. 2). While CD24 was expressed in most clusters, other stem-cell markers were enriched in distinct subsets: SOX9, CD44, and LGR4 expression marked clusters 3, 9, 10, and 11, whereas CD47 and EPCAM were highly expressed in clusters 5 and 8. The latter clusters were enriched for keratin (KRT)19, KRT7, and KRT8, indicating a cholangiocytic phenotype (Tarlow et al., 2014, MacParland et al., 2018). Cluster 9 expressed YAP1, MST1R, ETV4, and HNF4A, all associated with a hepatocytic fate (Fig. 2C) (Tarlow et al., 2014). These observations underscore the bipotent nature of EM organoids, with distinct stem cell populations primed towards either a cholangiocytic or a hepatocytic fate (Broutier et al., 2016, Huch et al., 2015).

When gene expression was analyzed by organoid donor, antigen presentation genes and ISGs were higher in HCV^+^ than NV organoids, with ISGs specifically upregulated in organoids from HCV1 and HCV3 donors. ISG expression was not observed in NV organoids but was induced after interferon α or β treatment (Fig. S2). Thus, the interferon response is intact in organoids derived from adult liver stem cells, and its constitutive activation in HCV1 and HCV3-infected organoids points to possible viral infection in these organoids.

### Stem cell organoids support long-term HCV replication

To test for a possible low-grade viral infection in the HCV^+^ organoids, we developed a droplet-digital RT-PCR assay to sensitively measure HCV RNA levels in cultures (Fig. 3A). HCV RNA was detected in HCV1 and 3, but not in NV or HCV2 organoids. RNAScope analysis detected HCV RNA in approximately 20% of cells for both HCV1 and HCV3 organoids (Fig. 3B). To ensure we detected active replication, we adapted the digital droplet PCR protocol to the negative strand of HCV RNA, an intermediate of viral replication (Klepper et al., 2017). HCV negative-strand RNA was detected in infected HCV1 and HCV3 EM organoids as well as in replication-competent Huh7.5 hepatoma cells infected with the HCV-Jc1 clone, confirming active RNA replication in these two samples of EM organoids (Fig. 3C).

**Figure 3.**
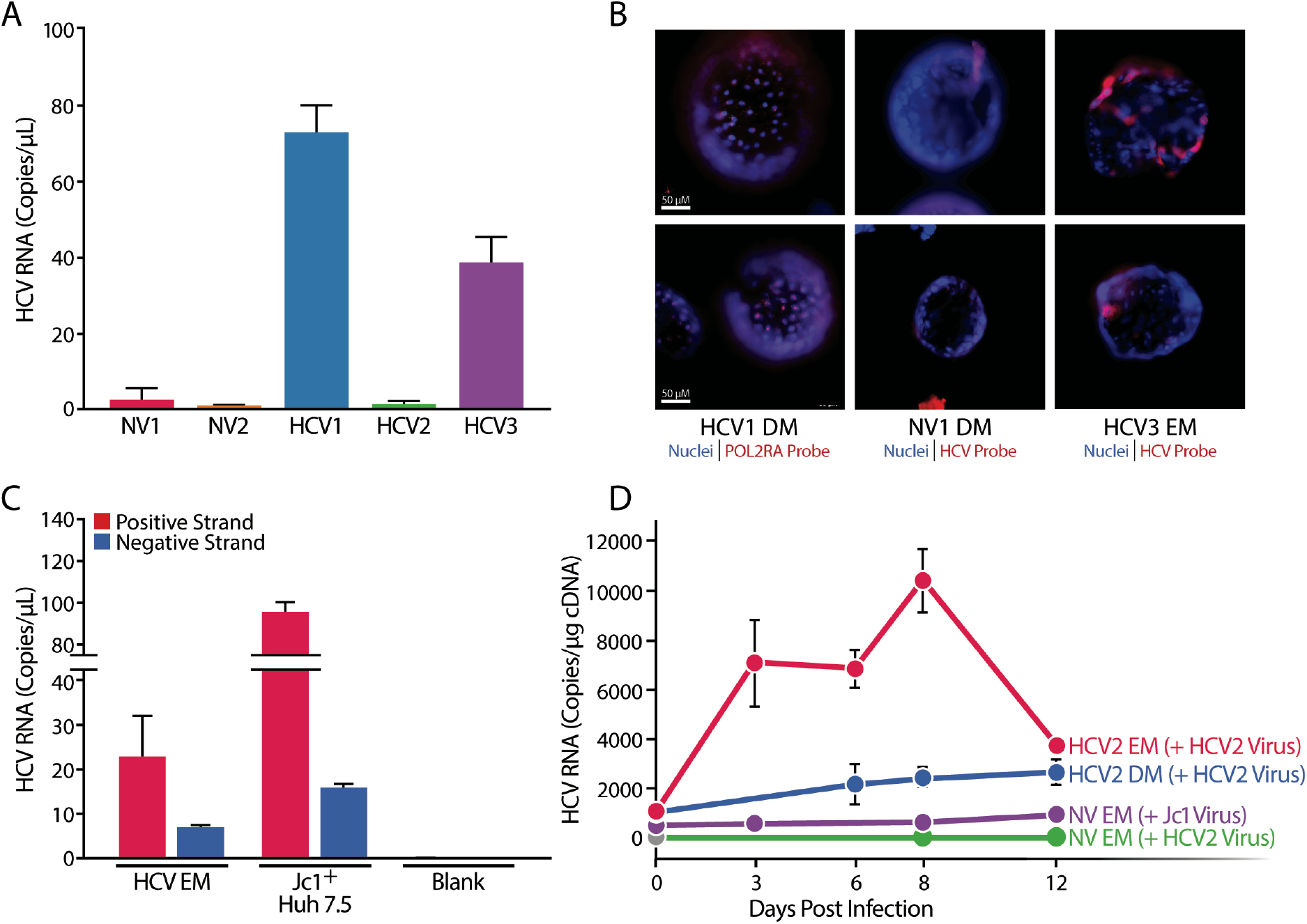
HCV infection persists in EM organoids from viremic donors. **(A)** ddPCR quantification of HCV RNA in EM organoids using a primer and probe set unique to the HCV 5’ UTR. Data are from n≥3 biological replicates per organoid donor and represented as mean ± SEM. **(B)** RNAScope of organoids using POL2RA or HCV RNA probes (red). Nuclei were stained with Hoescht (blue). **(C)** ddPCR quantification of positive and negative HCV strands in HCV1 organoids and HCV-infected Huh7.5 cells. Data are from n≥3 biological replicates per organoid donor and represented as mean ± SEM. **(D)** qPCR quantification of HCV RNA from EM and DM HCV2 organoids infected with autologous virus from the HCV2 donor and from NV EM organoids infected with virus from HCV2 or Jc1, over 12 days. EM organoids were passaged at day 8, resulting in a lower HCV concentration at day 12. In parallel, EM organoids from non-viral donors were spin-infected with the HCV2 viral isolate at the same MOI and with a JC1 lab strain at MOI = 5. Data are represented as mean ± SD.

Next, we focused on the HCV2 organoids, where no infection was detected (Fig. 3A). We utilized serum samples from the HCV2-infected individual to inoculate HCV2 EM and DM cultures by spin-infection (Fig. 3D). In both cultures, viral RNA was detected by quantitative RT-PCR over 2 weeks; EM cultures were more robustly infected than DM cultures. Notably, the same inoculum did not infect EM organoids from NV donors (Fig. 3D), highlighting a possible virus:host adaptation that may restrict infectivity in primary cell culture. These data show that liver stem cells from HCV^+^ donors are either infected or permissive to *ex vivo* infection with an autologous viral isolate.

### HCV infection drives transcriptional clustering in virus-inclusive scRNA-seq

HCV RNA is not poly-adenylated, and hence not captured by the standard poly-T capture oligonucleotide in the 10X Genomics Drop-seq protocol (Fig. 2). To distinguish the transcriptional profile of infected vs. uninfected cells, we designed a viral capture assay using a synthetic oligonucleotide corresponding to a conserved region of the HCV sequence in the Chromium Single-Cell 5’ Gel Bead & Library Kit (Wakita et al., 2005, Steinmann and Pietschmann, 2013, Catanese and Dorner, 2015). The protocol worked in Huh7.5 cells infected with HCV-Jc1 (Fig. S3). Next, we applied viral capture scRNA-seq to HCV1 EM cells under standard EM culture conditions or undergoing differentiation for 3 or 6 days. scVI analysis of the combined datasets identified 13 clusters (Fig. 4B, S4B, Data S2). Average HCV RNA expression was similar across the three differentiation stages, with an average of fewer than 10 copies per cell (Fig. S4). Cells with the highest HCV load (HCV^high^ cells) were in just three of the 13 clusters (Fig. 4A-C): cluster 4, which comprised infected EM and day-3 DM cells; and clusters 8 and 9, which comprised infected day-6 DM cells. These data indicate that HCV infection significantly perturbs gene expression in liver stem cells at their basal state and during differentiation.

**Figure 4.**
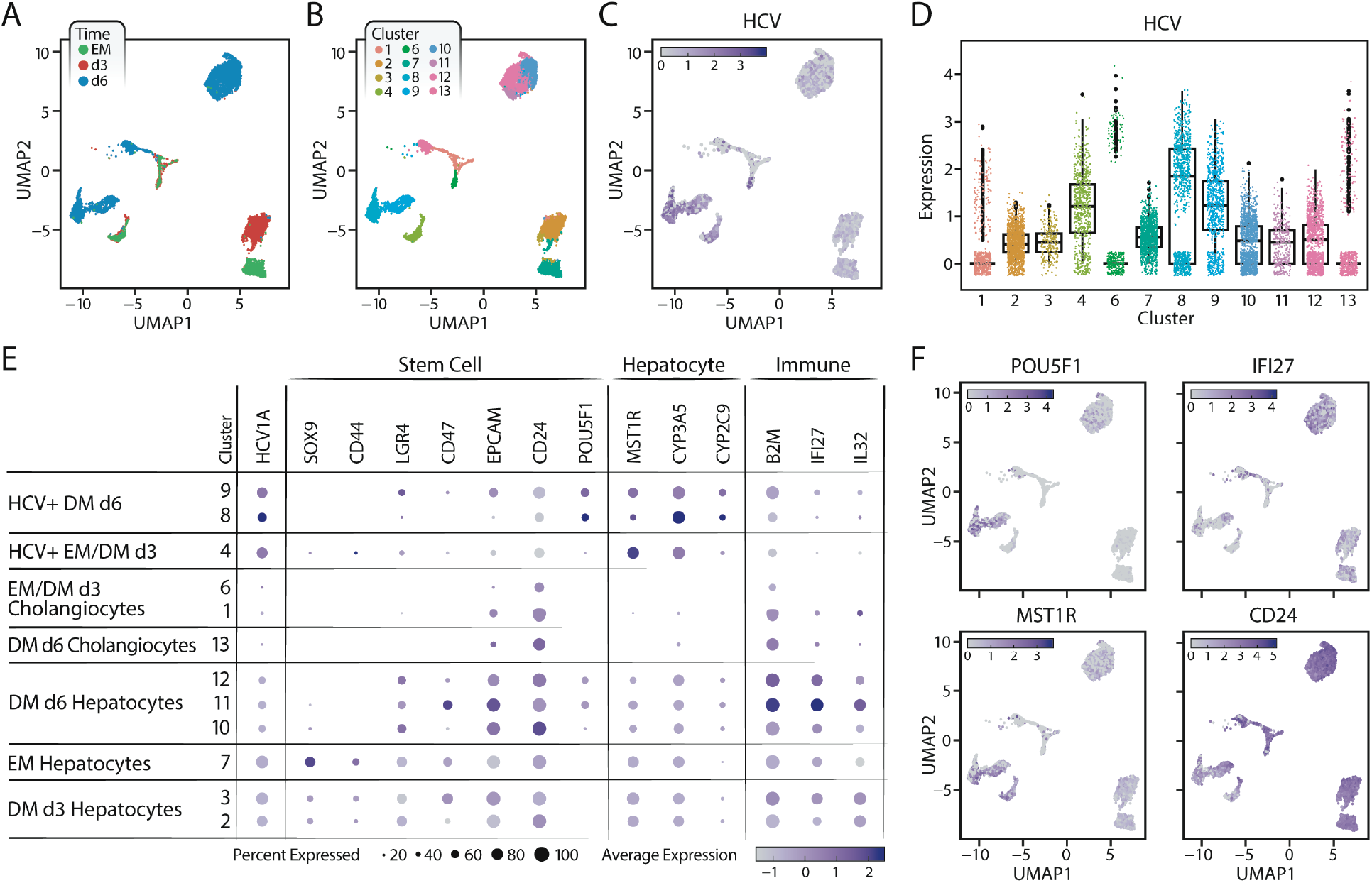
HCV infection of liver stem cells enhances differentiation to a hepatocyte-like fate. **(A)** A UMAP of HCV1 EM, day 3 (d3) DM, and day 6 (d6) DM organoids is generated from the 5’ scRNA-seq data with HCV capture oligo and plotted with cells identified by sample. **(B)** The same UMAP in (A) is plotted with cells identified by the 13 clusters from scVI analysis. Cluster 5 is omitted due to low read quality. **(C)** HCV RNA expression is plotted (as pseudolog) on the UMAP. **(D)** A box plot (as pseudolog) shows HCV RNA expression by cluster. **(E)** Dot plot shows gene expression (as pseudolog) of HCV (Viral), stem cell (Stem), hepatocyte, and immune markers. Clusters are on the y-axis, and are further labeled with identities assigned from gene expression analysis. **(F)** UMAPs are shown for gene expression (as pseudolog) of POU5F1, CD24, MST1R, and IFI27.

Uninfected or HCV^low^ cholangiocytic and hepatocytic progenitors were enriched in clusters 7, 2 and 3. EM cells (cluster 7) highly expressed hepatocyte progenitor markers, such as SOX9 and HNF4A (Chaudhari et al., 2016), TACSTD2 (TROP2), a gene expressed in bipotent liver progenitor cells, and CXCL8, a gene expressed in progenitor cells primed towards cholangiocyte differentiation (Aizarani et al., 2019) (Fig. 4E, F, S4D, E). On day 3 of differentiation, SOX9, CD44, TACSTD2 and CXCL8 were downregulated (clusters 2 and 3), but hepatocyte markers, such as GLUL and HAMP, were not upregulated (Aizarani et al., 2019, MacParland et al., 2018) (Fig. 4E, S4D). Upregulation of GLUL and HAMP occurred in uninfected day-6 DM cells (clusters 10–13), indicating these clusters include the most mature hepatocyte-like cells (Fig S4D).

Cholangiocyte-like cells (clusters 1, 6, and part of cluster 13) had low expression of stem cell and hepatocyte markers but high expression of cholangiocyte markers, such as KRT/8/19 (Fig. 4E, S4D). Notably, these clusters showed the lowest HCV infection rate, indicating that a cholangiocytic state does not support HCV infection (Fig. 4C, D, E).

### HCV infection perturbs markers of cellular differentiation and antiviral responses

We next compared the differentiation status of HCV^high^ clusters (4, 8, and 9) to that of lowly-infected clusters. HCV^high^ clusters showed low expression of hepatic stem cell markers, such as SOX9, CD44, LGR4, CD47, EpCAM, and CD24, at all stages of differentiation, including in the EM state (Fig. 4E). In contrast, MST1R, CYP3A5 and CYP2C9 hepatocyte markers were strongly expressed in all HCV^high^ clusters (Fig. 4E, F). These results indicate that HCV may perturb the stemness of progenitor cells to preferentially prime them toward a hepatocyte fate.

To exclude the possibility that clusters 4, 8 and 9 represent a subset of cells that occur during normal differentiation and are more permissive to HCV infection, we examined differentiated organoids from NV donors. scRNA-seq analysis of two NV organoids identified 14 unique clusters of cells (Fig S5A). However, none showed an expression pattern matching that of HCV^high^ clusters (Fig S5C), supporting the model that HCV infection actively perturbs stem-cell differentiation.

One notable difference between HCV^high^ and HCV^low^ clusters was that upon differentiation, HCV^high^ cells in clusters 8 and 9 upregulated expression of POU5F1 (OCT4) (Fig. 4F), a master regulatory transcription factor defining embryonic stem cells and usually absent from liver progenitor cells or lowly expressed during differentiation (Simandi et al., 2016). When upregulated in tissue progenitor cells, OCT4 promotes a cancer stem-cell state (Zhu et al., 2015). These data support a model where HCV infection of liver stem cells dysregulates differentiation into hepatocyte-like cells with a possible cancer stem cell identity.

We further compared expression of interferon-stimulated and antigen presentation genes between HCV^low^ and HCV^high^ clusters -including IFI27, IFI6, and B2M (Fig. 4E, F). In stem cells, expression of all three genes was lower in HCV^high^ compared to HCV^low^ clusters,. Expression increased during differentiation of HCV^low^ cells and was highest in cluster 11 comprised of mostly uninfected DM d6 hepatocytes (Fig. 4E). However, in HCV^high^ clusters expression did not increase on differentiation. These results show that interferon-stimulated and antigen presentation genes are induced in uninfected bystander cells at all stages of differentiation while in HCV-infected cells this response is effectively blunted. They underscore results from previous studies reporting increased interferon responses upon cellular differentiation (Wu et al., 2018).

### Network propagation identifies alterations in splicing, ATP synthesis and ribosome biogenesis pathways in HCV^hlgh^ clusters

To identify the main transcriptional targets of HCV infection, we compared the top 150 up- or down-regulated genes from each HCV^high^ cluster. We found 81 that were up-regulated and 142 down-regulated within at least two clusters, relative to uninfected clusters (Fig. S6A and B). Based on gene ontology term enrichment analysis, upregulated pathways included RNA splicing, extracellular matrix organization and cell morphogenesis involved in differentiation (Fig. 5A, S6), and down-regulated pathways included ribosomal genes and oxidative phosphorylation (Fig. 5B, S6) (Zhou et al., 2019).

**Figure 5.**
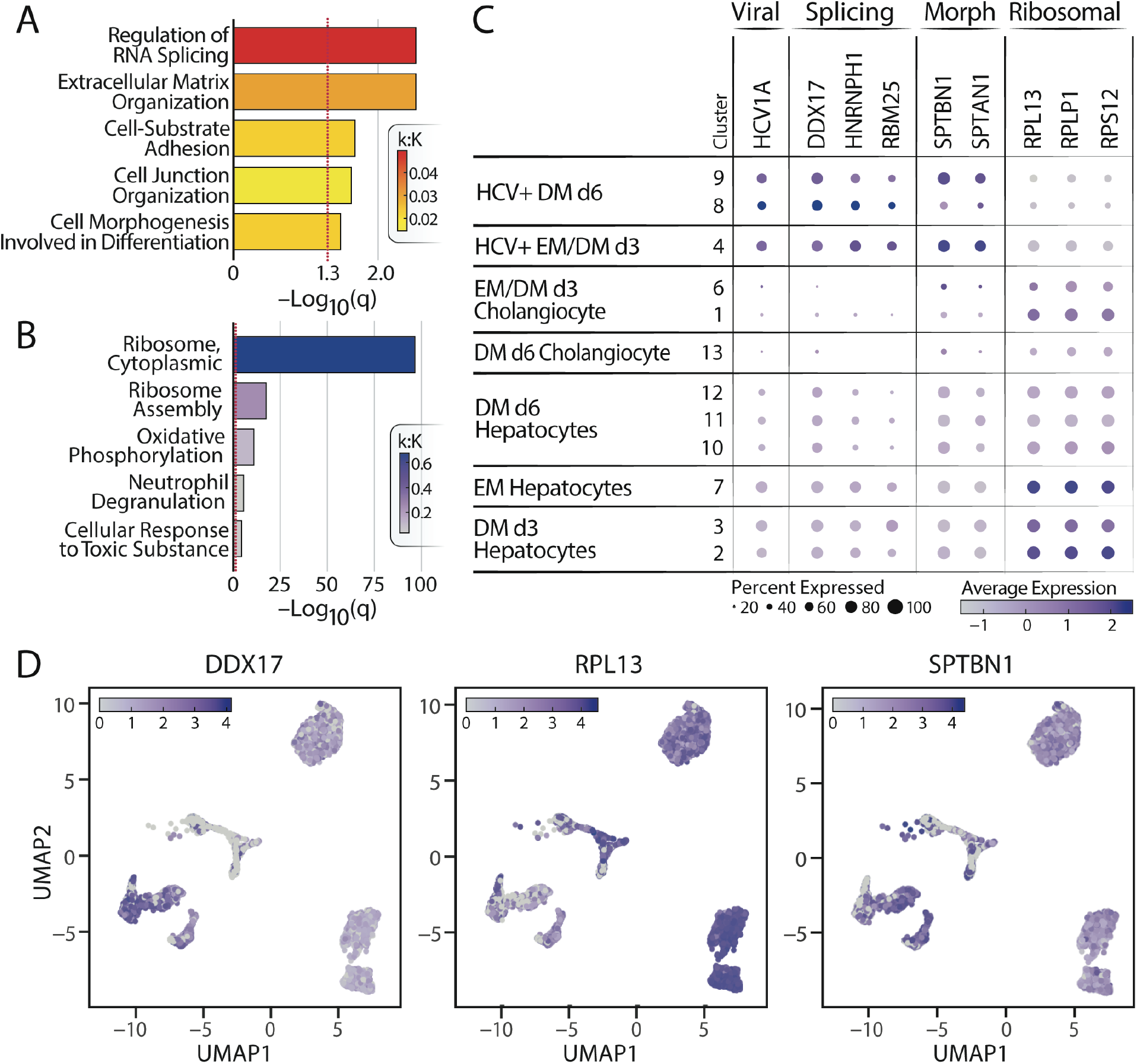
HCV infection of liver stem cells upregulates pro-viral factors and RNA splicing genes. Metascape gene ontology term enrichment analysis identifies pathways from the 81 genes commonly up-regulated **(A**) or down-regulated **(B)** in at least two of the HCV^high^ (4, 8 and 9) clusters. Terms are ordered by q-value and colored based on the observed genes within the term (k) divided by the total genes in the term (K). **(C)** A dot plot is shown for genes in different GO terms: RNA splicing (splicing), cell morphogenesis involved in differentiation (morph), and ribosome. Clusters on on the y-axis and labeled as in Fig. 4. **(D)** UMAPs show expression (as pseudolog) of three of the top dysregulated genes: DDX17, RPL13, and SPTBN1.

We noted that many up- or downregulated genes were also detected in a previously published HCV protein interactome in Huh 7.5 cells (Ramage et al., 2015). In particular, a network propagation analysis between the commonly differentially expressed genes in the scRNA-seq and the HCV protein interaction partners revealed three major converging molecular networks: ribosome biogenesis and mitochondrial transport/ATP synthesis, comprised of mostly down-regulated genes, and splicing, composed of mostly up-regulated genes (Fig 6A). This observation indicates that viral infection intersects these cellular pathways at multiple steps.

**Figure 6.**
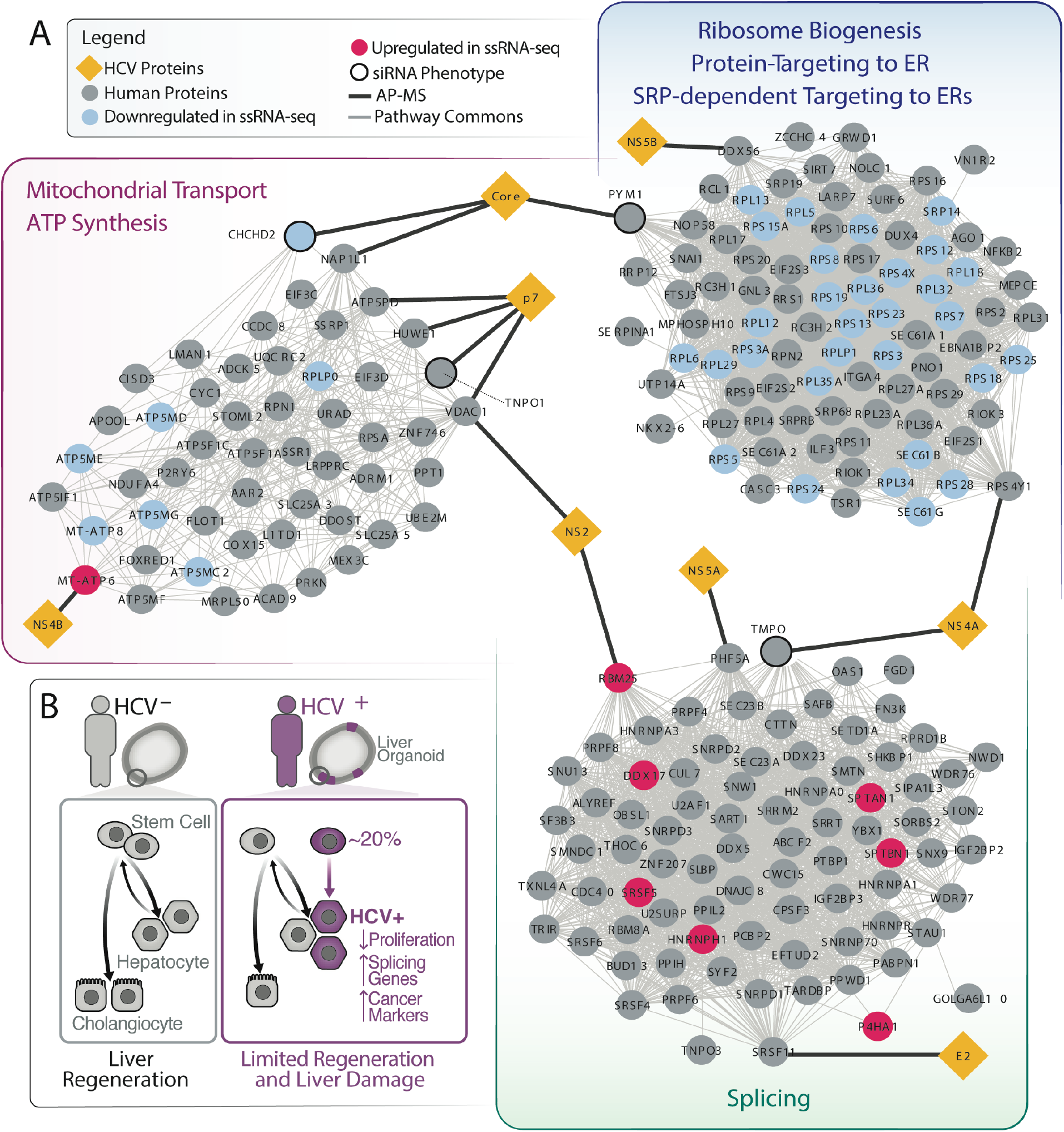
Genes regulated by HCV infection of liver stem cells form networks with HCV interacting proteins. **(A)** Network analysis of commonly up- or down-regulated genes from 5A and 5B and previously published HCV interactome. **(B)** Model of HCV infection in liver stem cells.

Down-regulation of pathways involved in ribosome biogenesis, mitochondrial transport/ATP synthesis and oxidative phosphorylation suggest that HCV^high^ cells have reduced proliferation rates (Gerresheim et al., 2019) (Fig. 5, Fig. 6, Data S3). Up-regulated pathways such as splicing and cell morphogenesis support the notion that HCV positive cells have altered differentiation (Sen et al., 2013, Bhate et al., 2015, Hyun et al., 2020, Yamazaki et al., 2018, Grammatikakis et al., 2016, Dardenne et al., 2014).

In summary, extensive transcriptional reprogramming is observed in infected organoid cells that skews differentiation, reduces proliferation, antagonizes interferon signaling, promotes tumorigenesis and supports viral replication.

## Discussion

Stem cells have so far not been considered a significant reservoir for HCV infection in the liver given reports of high constitutive levels of ISG expression in iPSCs and the difficulty in obtaining culturable material (Marukian et al., 2011, Ploss et al., 2010). By generating adult stem cell-derived organoids from donors that were actively replicating HCV at the time of liver resection and keeping these organoids for months in culture, we demonstrate that liver stem cells represent novel, long-lived HCV-producing cells *ex vivo* and possibly *in vivo*. Notably, we showed that the three predominant genotypes of HCV, HCV 1, 2 and 3, can each replicate in liver stem cells.

By performing virus-inclusive scRNA-Seq of infected and differentiating stem cell organoids, we find characteristic differences between HCV^+^ and HCV^-^ cells. 1) HCV^high^ clusters lack ISG expression, consistent with the notion that HCV infection effectively antagonizes interferon signaling, i.e. through its core and NS3/4 proteins (Wong and Chen, 2016). However, HCV modulates the interferon response only in infected, but not in uninfected, stem cells and hepatocyte-like cells, limiting viral spread to bystander cells with active interferon responses. Overall, adult stem cell-derived organoids showed intact interferon responses to exogenous interferon treatment and viral infection, albeit lower in EM cultures than in DM cultures, possibly explaining increased infection by HCV in stem cell organoids. Constitutive high ISG expression was not observed in adult stem cell-derived organoids. Interestingly, EM organoids harboring HCV-1 and HCV-3 showed different patterns of ISG and MHC gene expression. These differences could reflect the differing viral genotypes or donor genetic backgrounds, which will both be further explored using the organoid platform.

2) Aside from silencing ISGs, HCV infection of stem cells perturbs the expression of differentiation genes. For instance, HCV^high^ cells up-regulate hepatocyte markers including CYP2C9, which was previously shown to be higher in HCV^+^ versus HCV^−^ non-tumoral liver tissue (Higgs et al., 2013). Further, HCV^high^ cells upregulate several genes that influence cell differentiation including SPTNB1, a gene which negatively regulates proliferation of quiescent hepatocytes after partial hepatectomy (Thenappan et al., 2011, Tang et al., 2003), and numerous genes for RNA splicing. Notably, alteration of splicing in adult mouse livers led to impaired hepatocyte maturation (Bhate et al., 2015), supporting a model where HCV infection perturbs proper liver regeneration by manipulating RNA splicing. Splicing factors DDX5, DDX17, and HNRNPH1 facilitate cell-fate determination (Dardenne et al., 2014, Grammatikakis et al., 2016, Yamazaki et al., 2018) and prevent the epithelial-mesenchymal transition when expressed at high levels (Dardenne et al., 2014). RNA splicing proteins also interact with several HCV proteins. During HCV infection, DDX5 and HNRNPH1 are translocated from the nucleus to the cytoplasm by HCV NS5B or Core proteins to promote viral replication (Kuroki et al., 2013, Goh et al., 2004, Lee et al., 2011). Our results suggest that HCV infection upregulates these factors to promote its replication while also skewing differentiation towards a hepatocytic fate.

3) At the same time, HCV^high^ cells strongly down-regulate stem-cell markers except for POU5F1 (OCT4). HCV core protein regulates expression of OCT4 in HCV^+^ HCC cells, and increased OCT4 expression drives cell-cycle progression (Zhou et al., 2016). The down-regulation of normal liver stem-cell factors with upregulated OCT4 may allow HCV-infected stem cells to replicate without maintaining the normal regenerative properties of hepatocytes, which would facilitate the development of cancer stem cells.

Combined, our observations indicate that HCV^+^ hepatocyte-like cells have reduced proliferative and trans-differentiation capacities, which could hamper their ability to regenerate liver tissue and increase the potential for liver damage in chronically-infected patients. Our gene expression findings suggest a model in which infection with HCV alters the differentiation of bi-potent liver stem cells by dampening cellular proliferation and mitochondrial function while upregulating cellular splicing, hepatocyte markers and pluripotent stem cell factor OCT4 (Fig 6B).

Our study is limited by small sample size and low viral replication in the organoids. Since most HCV patients are now treated with direct-acting antivirals, which usually eradicate the virus, access to liver tissue from donors carrying active HCV is limited. Moreover, HCV replicates at approximately 50 to 100-fold lower levels in liver organoids as compared to transformed Huh7.5 hepatoma cells. Therefore, ultrasensitive assays including ddPCR, RNA-Scope and virus-inclusive scRNA-seq are necessary to track HCV replication in organoids, but only allow limited studies of the viral life cycle. Future studies will explore whether stem cell reprogramming is dependent on active viral replication or may persist beyond viral clearance, which would explain lasting liver damage and altered cancer risk in individuals freed of the virus.

## Acknowledgments

We thank the Gladstone Genomics Core and CZ Biohub Genomics platform for help with library preparation and sequencing, and the Gladstone Microscopy Core for help with microscopy. We thank Gorica Margulis for lab training and access to the CZ Biohub Genomics platform and Spyros Darminis for help with sequencing at the CZ BioHub. We thank Tiffany Kanegawa from 10x Genomics on design of scRNA-seq with custom oligo for HCV detection. We acknowledge Dr. Charles Rice at Rockefeller University for providing Huh 7.5 cells. We thank Holger Willenbring, Jackie Maher, and the UCSF Liver Center for funding, resources, and advice. We thank the Ibrahim El-Hefni Liver Biorepository at the California Pacific Medical Center Research Institute for providing patient tissue samples used to generate liver organoids. We thank Amy Kistler, Laurent Coscoy, Peter Sarnow, and Karla Kirkegaard for helpful discussions to inform experimental design and guide data analysis.

## Author contributions

Conceptualization, N.L.M. and M.O.; Methodology and Investigation, N.L.M., C.R.S., A.L.E., M.M.K., T.Y.T, F. Z. and V.N.; Writing – Original Draft, N.L.M., D.E.L. and M.O.; Software and Formal Analysis, N.L.M., T.A., D.E.L., M.B., and T.T.N.; Visualization, N.L.M. and D.E.L.; Supervision, J.L.B., A.K., N.N., S.R.Q., T. M., N.J.K., S.C., T.C.M., N.Y., and M.O.; Funding Acquisition, M.O.

## Data and materials availability

All of the data that is underlying to this work will be deposited in the public GEO database upon acceptance of the manuscript for publication.

## STAR Methods

### KEY RESOURCES TABLE

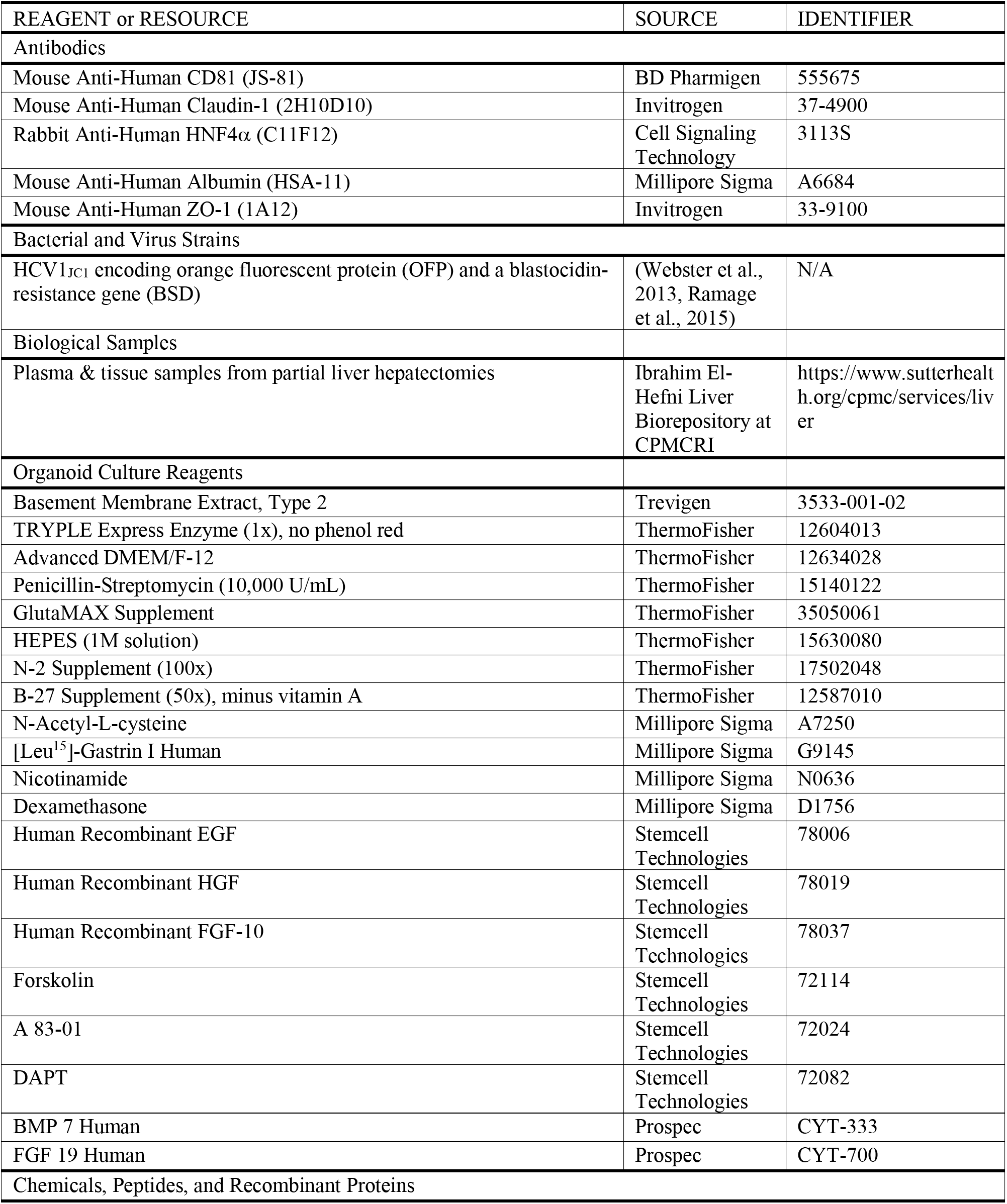
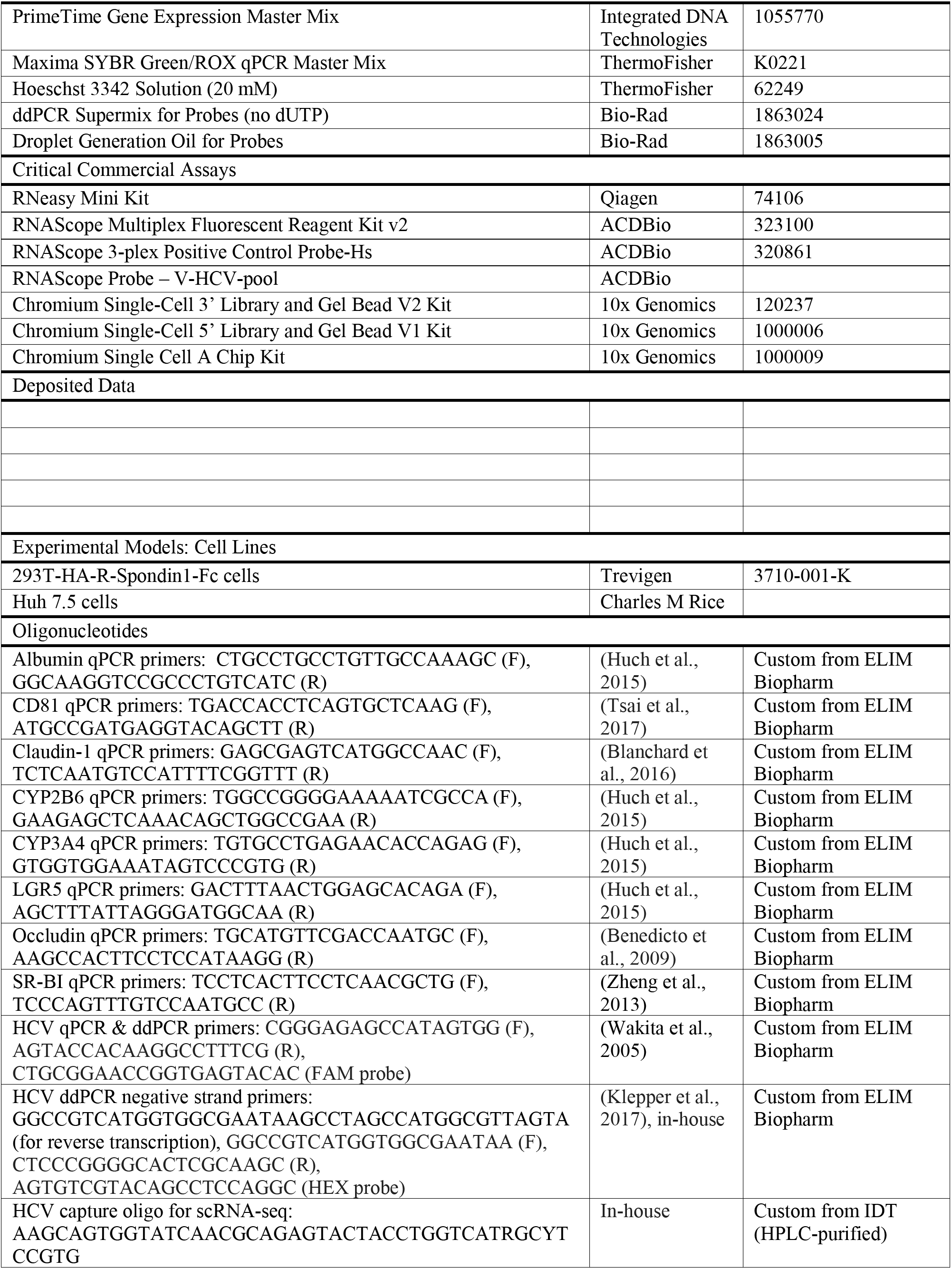
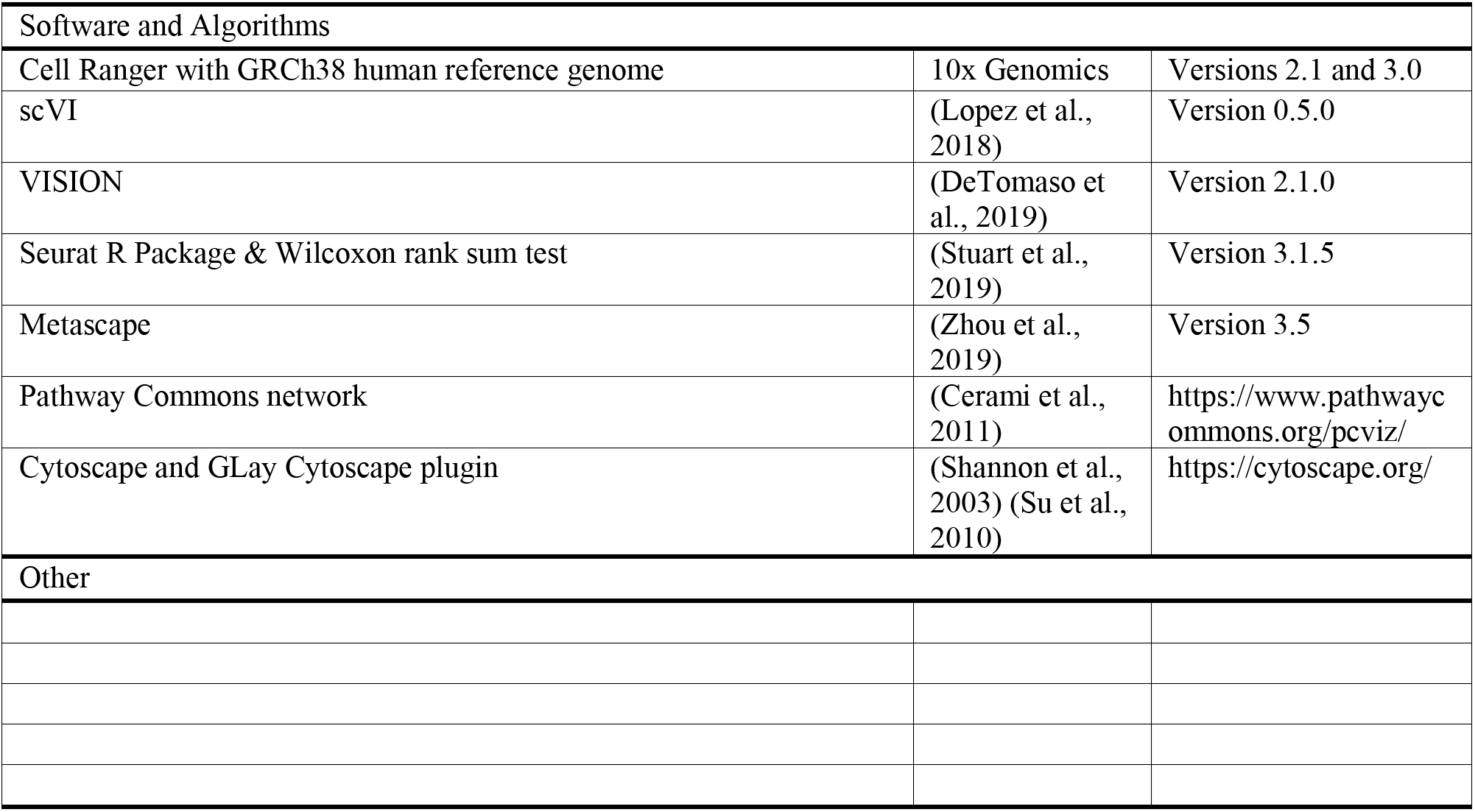

### CULTURE OF CELL LINES

293T-HA-R-Spondin1-Fc cells were purchased from Trevigen (Catalogue Number 3710-001-K) and cultured according to the manufacturer’s protocol to generate conditioned medium of R-spondi-1. Briefly, cells were grown in selection growth medium (DMEM with 10% FBS, 1% penicillin-streptomycin, 1% glutamine, and 100 mg/mL Zeocin) for >5 days until they were >90% confluent. Medium was replaced with organoid basal medium (Advanced DMEM/F12 from Invitrogen supplemented with 1% penicillin-streptomycin, 1% Glutamax, and 10 mM HEPES). After 3 days, cell supernatant (i.e., R-spondin-1 conditioned medium) was collected, centrifuged at 3,000 x g for 15 min, filtered through a 0.22-μm filter, and frozen at −20°C in 10 mL aliquots. This process was repeated by adding fresh organoid basal medium to the cells and collecting supernatant after 4 days.

Huh 7.5 cells were provided by Charles M. Rice and grown under standard conditions.

### CULTURE OF HUMAN LIVER ORGANOIDS

Liver organoids were generated from bi-potent liver stem cells as described (Broutier et al., 2016, Huch et al., 2015). Briefly, single cells were isolated from the healthy resection margins of liver samples obtained during partial hepatectomy. After tissue digest, the heterogeneous mixture of single cells was either directly plated or further enriched for EPCAM+ cells using the EasySep™ Human EpCAM Positive Selection Kit and then plated. For plating, cells were suspended in cold organoid basal medium and mixed with two parts reduced growth factor BME2 (Basement Membrane Extract, Type 2, Trevigen, catalogue number 3533-001-02). From this mixture, 50-μL drops containing 1,000–20,000 cells were seeded in 24-well suspension culture plates (GreinerBio-one, catalogue number 662-102). Drops were incubated at 37°C for >20 min and solidified. After this, 500 μL of expansion medium (EM) was added to each well. EM is organoid basal medium supplemented with 1% N_2_ and 1% B27 (both from Gibco), 1 mM N-acetylcysteine (Millipore Sigma), 10 nM [leu^15^]-gastrin I human (Millipore Sigma), 10% (vol/vol) R-spondin1 conditioned medium, and 10 mM nicotinamide (Millipore Sigma). EM additionally contains 50 ng/ml recombinant human EGF, 25 ng/ml recombinant human HGF, 100 ng/ml recombinant human FGF10, 10 μM Forskolin, and 5 μM A83-01 (all from Stem Cell Technologies). EM was replaced every 3–4 days.

After 14–21 days, organoids were passaged. For this, cold basal medium was used to collect organoids in 15-ml Falcon tubes and dissolve BME2. Tubes were centrifuged at 250 x g for 5 min at 4 °C. Medium was aspirated, TRYPLE (Gibco) was added to the organoids, and the mixture was incubated at 37°C for 5-10 min. Organoids were further dissociated by pipette mixing and then diluted in cold basal medium. After another spin and medium aspiration, cells were mixed with BME2 and seeded into new drops. After this initial passage, organoids were passaged every 2 weeks. Stocks of early-passage (P1 to P3) organoid lines were prepared by dissociating organoids, mixing them with recovery cell culture freezing medium (Gibco), and frozen, following standard procedures. These samples could be thawed and immediately cultured, using EM supplemented with 10 μM Y-27632 (Stem Cell Technologies) for the first 3 days after thawing.

To induce differentiation to a hepatocyte-like fate, EM was supplemented with 25 ng/ml BMP7 (ProSpec) for 3–4 days and then changed to differentiation medium (DM). DM is basal medium supplemented with 1% N_2_, 1% B27, 1 mM N-acetylcysteine, 10 nM [leu^15^]-gastrin I human, 50 ng/ml EGF, 25 ng/ml HGF, 0.5 μM A83-01, 25 ng/ml BMP7, 10 μM DAPT (Stem Cell Technologies), 3 μM dexamethasone (Millipore Sigma), and 100 ng/ml recombinant human FGF19 (ProSpec). DM was changed every 3–4 days for a period of 3–15 days.

### REAL TIME QUANTITATIVE PCR

RNeasy kits from Qiagen were used for RNA extraction and isolation. To extract RNA, medium was aspirated from the well to leave organoids suspended in BME2. 350 μL of buffer RLT (lysis buffer) was added directly to the well. After a short incubation for RLT to dissolve BME2, sample lysate was transferred to a 1.5-mL Eppendorf tube. Lysate was either stored at −80°C for up to 1 month or extracted immediately, following the manufacturer’s protocol for the RNeasy kit. Final RNA concentrations were measured with a NanoDrop ND-1000. Total RNA was reverse-transcribed using oligo(dT)_18_ primers (Thermo Scientific), random hexamers primers (Thermo Scientific), and AMV reverse-transcriptase (Promega). cDNA was diluted to 5 ng/μL. Gene expression was assayed by real-time quantitative PCR using Maxima SYBR Green qPCR Master Mix (Thermo Scientific) on an ABI 7900HT real-time PCR system. The SYBR Green qPCR reactions contained 10 μL of 2x SYBR Green Master Mix, 2 μL of diluted cDNA, and 8 pmol of both forward and reverse primers. The reactions were run using the following conditions: 50°C for 2 min, 95°C for 10 mins, followed by 40 cycles of 95°C for 5 secs and 60°C for 30 secs. Relative values for each transcript were normalized to 18S rRNA. Gene primers used are listed in Key Resources. For every qPCR run, three technical replicates per sample were used for each gene.

For HCV detection, we used HCV-specific primers and a FAM-conjugated Taqman probe (Applied Biosciences) with PrimeTime Gene Expression Master Mix (IDT) as described (Wakita et al., 2005). Sequences were as follows: 5’-CGGGAGAGCCATAGTGG-3’ (forward), 5 ‘-AGTACCACAAGGCCTTTCG-3’ (reverse), 5 ‘-CTGCGGAACCGGTGAGTACAC-3’ (probe). For quantification of viral copies, an HCV standard was generated by serial dilution of a described JCV1-mKO2 plasmid (Webster et al., 2013) and used in every qPCR run.

### DROPLET-DIGITAL (dd) PCR TO DETECT HCV

We used Bio-Rad’s QX100 Droplet Digital PCR (ddPCR) System to detect HCV at low levels in the organoids. Organoid RNA was extracted and reverse-transcribed to cDNA as described above. For each sample, 40 ng of cDNA was mixed with 10 μL of 2x ddPCR Supermix for Probes (no dUTP) (Bio-Rad), 18 pmol of forward and reverse primers for HCV (described above), and 4.5 pmol of the HCV FAM probe (described above). This mixture was dispensed into the sample wells of a D8 droplet generator cartridge (Bio-Rad), and droplets were generated using the QX100 droplet generator according to the manufacturer’s instruction. After droplet generation, the reaction mix was transferred to a ddPCR 96-well PCR plate (Bio-Rad), and the plate was sealed. The plate was run on a thermal cycler with the following conditions: 95°C for 10 min, 45x cycles of 94°C for 30 sec and 59.4°C for 1 min, 98°C for 10 min, and 4°C for 30 min. After the protocol was completed, the plate was transferred to a QX100 droplet reader for analysis. For every ddPCR run, two technical replicates per sample were used. A water blank and positive control were included in every run. The positive control was cDNA from Huh 7.5 s infected at MOI = 0.01 with the JC1-mKO2 HCV strain mentioned above.

To detect the negative strand of HCV, we adapted a qPCR assay for ddPCR (Klepper et al., 2017). In our assay, RNA is extracted from organoids and reverse transcription performed with a tagged forward primer (Tag-RC1) and ThermoScript Reverse Transcriptase (Thermo Scientific). The Tag-RC1 sequence is 5’-GGCCGTCATGGTGGCGAATAAGCCTAGCCATGGCGTTAGTA-3’. For first strand synthesis, RNA was mixed with Tag-RC1 primer and dNTP and the whole mix was incubated at 70°C for 8 min then 4 °C for 5 min. For reverse transcription, ThermoScript Reverse Transcriptase, enzyme buffer, DTT and RNAse inhibitor were added to the reaction. The total mix was incubated at 60°C for 1 hr then heated to 85°C for 5 min to terminate the reaction. To degrade input RNA, RNAse H was added to the complete reaction followed by incubation at 37°C for 20 min. After this cDNA was analyzed by ddPCR as described above but with different primers and probe. We used the described tag-specific forward primer (5’-GGCCGTCATGGTGGCGAATAA-3’) and reverse primer RC21 (5’-CTCCCGGGGCACTCGCAAGC-3’) (Klepper et al., 2017). We designed a HEX-conjugated Taqman probe (Applied Biosciences) compatible with these primers with sequence 5’-AGTGTCGTACAGCCTCCAGGC-3’.

### WHOLE-MOUNT ORGANOID STAINING

Organoids were processed for imaging as previously described (Broutier et al., 2016). Briefly, organoids were removed from BME2 with 3x cold PBS washes then fixed in 2-3% paraformaldehyde for 30-60 min and washed 3x in PBS. Fixed organoid samples were stored at 4°C for up to 2 months.

For staining, organoids were blocked in PBS supplemented with 0.5% Triton X-100, 1% DMSO, 1% BSA, and 1% donkey or goat serum. Organoids were blocked overnight at 4°C. Blocking solution was then removed and replaced with blocking solution containing primary antibodies diluted 1:250. Organoids were incubated with primary antibodies for 48 hr at 4°C. After this, organoids were washed 3x in PBS and incubated overnight at 4 4°C with secondary antibodies diluted 1:250 in PBS. Organoids were washed 3x in PBS and stained with Hoescht before visualization. Organoids were imaged on a Zeiss Lightsheet Z.1. Images were processed using a combination of the Zeiss software, ImageJ 1.51f, and Imaris 9.3. Primary antibodies we used include CD81 (BD Pharmigen, JS-81), Claudin-1 (ThermoScientific, 2H10D10), HNF4□ (Cell Signaling Technology, C11F12), Albumin (Sigma-Aldrich, HSA-11), and ZO1 (ThermoScientific, 1A12).

#### RNAScope

For RNAScope, organoids were fixed as described above, except incubation in 2% paraformaldehyde was extended overnight. The RNAScope kit, HCV probe and 3-plex positive control probe were purchased from ACDBio. To process and stain fixed organoids, we followed the manufacturer’s instructions. Organoids were imaged on a Zeiss Axio Observer Z. 1 and Zeiss Lightsheet Z.1. Images were processed using a combination of the Zeiss software, ImageJ 1.51f, and Imaris 9.3.

### HCV INFECTION OF HUH 7.5 CELLS

Huh 7.5 cells were infected with a previously described monocistronic infectious clone of HCV1_JC1_ encoding orange fluorescent protein (OFP) and a blastocidin-resistance gene (BSD) (Webster et al., 2013, Ramage et al., 2015). Cells were infected at MOIs of 0.01, 0.1, and 0.2. Seven days later, cells were processed for single-cell RNA-sequencing as described below.

### HCV INFECTION OF ORGANOIDS

HCV2 EM and DM day 3 organoids were spin-infected with patient-matched virus as follows. Organoids were collected and lightly dissociated by a 3 min incubation at 37°C with TRYPLE. Cells were then mixed with sera (HCV titer of 3.5e6 IU/mL) at an MOI of 450 IU/organoid in a 24-well suspension culture plate. The plate was centrifuged at 600 x g for 1 hr at room temperature, followed by a 2 hr incubation at 37°C. After this, cells were collected and washed 3x in basal media. Washed cells were centrifuged at 250 x g at 4°C for 5 min and then seeded in fresh BME2 drops in a new 24-well suspension culture plate. Every 2-3 days, RNA was harvested as described above. On day 9 post-infection, EM organoids were thinly passaged.

### SINGLE-CELL RNA-SEQUENCING

Single-cell RNA sequencing was performed using the 10x Genomics Chromium System (Zheng et al., 2017). EM and DM organoids samples were processed within 1 month of starting the organoid culture, or within 3 weeks of thawing a frozen stock. To prepare single-cell suspensions for organoids, samples were incubated with TRYPLE for 10-15 min, washed once with basal media, and flowed through 40 μm cell strainers to remove cell aggregates. Cell and viability counts were performed using trypan blue and hemocytometers. Viability was >75% for all organoid samples. To prepare single-cell suspensions for Huh 7.5 cells, samples were incubated with 0.25% Trypsin-EDTA solution (Gibco) for 10-15 min, washed once with DMEM, and flowed through 40 μm cell strainers to remove cell aggregates. Viability was >94% for all Huh 7.5 samples. Single-cell suspensions were concentrated at 12,000 target cells in 30 μl of basal media, due to an estimated lower retention rate for hepatocytes per discussion with 10x Genomics.

Single-cell RNA-seq libraries were generated using either the Chromium Single-Cell 3’ Library and Gel Bead V2 Kit or the Chromium Single-Cell 5’ Library and Gel Bead V1 Kit following the manufacturer’s protocols. For the 5’ Library and Gel Bead Kit, we added a custom HCV capture oligo (5’-AAGCAGTGGTATCAACGCAGAGTACTACCTGGTCATRGCYTCCGTG-3’) to the master mix for reverse transcription at a 1:1 equimolar ratio with the Poly-dT RT Primer. We proceeded directly from cDNA Amplification & QC to 5’ Gene Expression (GEX) Library Construction.

Single-cell libraries were loaded on an Illumina HiSeq 4000 System or NovaSeq 6000 System with an S1 flow cell with the following reads: 26 bases Read 1 (cell barcode and unique molecular identifier (UMI), 8 bases i7 index 1 (sample index) and 98 bases Read 2 (transcript).

### SINGLE-CELL DATA ANALYSIS

The Cell Ranger pipeline (10X Genomics, Pleasanton, CA, USA) (Version 2.1 and 3.0) was used to perform sample de-multiplexing, barcode processing, and 3’ UMI counting. FASTQ files were aligned to the human reference data set (GRCh38) from 10x Genomics. When applicable, the human reference data set was appended with the HCV genome (GenBank ID AB047639.1 to align libraries from Huh 7.5 cells infected with a JFH1-derived strain and GenBank ID AF009606.1 to align libraries from HCV1 organoid samples of genotype 1).

For each experiment, filtered gene expression matrices from each sample were read into scVI (version 0.5.0) (Lopez et al., 2018) and jointly analyzed using default hyperparameters. Donor annotation was used as batch factor in all donor-derived samples, and library identifier was used as the batch factor in the Huh7.5 experiment. The 10-dimentional latent representation from scVI was then clustered, visualized, and explored using VISION (version 2.1.0) (DeTomaso et al., 2019). One low quality cluster from the analysis of 5’ scRNA-seq presented in Figure 4, Cluster 5, was removed from visualization.

Differentially expressed genes (DEGs) were determined for clusters using the Wilcoxon rank sum test implemented in Seurat R Package (version 3.1.5) (Stuart et al., 2019) with min.pct = 0.25. Sequencing statistics, including number of cells per sample, reads per cell, and genes per cell are described in Table S1.

### GENE ONTOLOGY TERM ENRICHMENT (GOTE) ANALYSIS

We performed GOTE analysis using Metascape (version 3.5) (Zhou et al., 2019). Express Analysis default settings were used.

### NETWORK PROPAGATION INTEGRATIVE ANALYSIS

We performed network propagation to integrate the single-cell RNA sequencing (scRNA-seq) data with the HCV virus-host protein-protein interactome using the Pathway Commons network (Cerami et al., 2011). Specifically, we used a heat-diffusion kernel analogous to random walk with restart (RWR, also known as insulated diffusion and personalized PageRank) which better captures the local topology of the interaction network compared to a general heat diffusion process. The process is captured by the steady-state solution as follows:

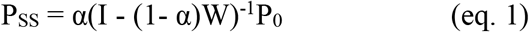

where P_SS_ represents the vector of propagated values at steady-state, P_0_ is the initial labeling (genes of interest from molecular studies), W is the normalized version of the adjacency matrix of the underlying network (in this implementation W = AD^-1^, where A is the unnormalized adjacency matrix, and D is the diagonal degree matrix of the network), I is the identity matrix, and α denotes the restart probability (here, α=0.2), which is the probability of returning to the previously visited node, thus controlling the spread through the network.

We performed two independent propagations, one for the scRNA-seq and one for the HCV interactome. For the scRNA-seq, we compiled all differentially expressed genes (up or down). For the HCV interactome, we used the 134 unique human preys from the published HCV interactome done in Huh7 cells, based on a HCV MIST score greater than 0.68 and passing the 99^th^ percentile for compPASS scoring (Ramage et al., 2015). To control for nodes with high degree (i.e. many connections), which due to their heightened connectivity are biased to receive higher propagation scores, we conducted a permutation test. Specifically, we simulated random propagations by shuffling the positive scores to random genes, repeating this 20,000 times for both propagation runs. Next, we calculated an empirical p-value by calculating the fraction of random propagation runs greater than or equal to the true propagation run for each gene. We integrated the data by extracting the genes with a p<=0.03 in both propagation runs.

The network was created by extracting the subnetwork of significant genes from the same Pathway Commons network visualizing interconnected genes using Cytoscape (Shannon et al., 2003). The resulting network was clustered into subnetworks using the GLay Cytoscape plugin (Su et al., 2010). One cluster was omitted from visualization in Figure 6 because it contained less than 5 genes, information about this cluster is included in Data S3.

## Supplementary Materials

Figure S1. Related to Figure 1. EM Organoids have Similar Lifespans and Viability between HCV Positive and Negative Donors.

Figure S2. Related to Figure 2. Liver Organoids are Responsive to Interferon Treatment.

Figure S3. Related to Figure 3 and Figure 4. HCV Infection of Hepatoma Cells

Figure S4. Related to Figure 4, 5, and 6. Uninfected cells differentiate into both hepatocyte and ductal like cells

Figure S5. Related to Figure 4. Differentiated cells from Non viral donors do not have clusters similar to HCV^high^ clusters

Figure S6. Related to Figure 5 and 6. HCV Infection of Liver Stem Cells Down-Regulates Oxidative Phosphorylation.

Table S1. Related to Figures 2, 4, and 5. Single-Cell RNA-sequencing Statistics for Liver Organoid Samples.

Table SI1: Related to Figure 2. Differentially expressed genes by cluster from 3’ scRNA-seq.

Table SI2: Related to Figure 4 and 5. Differentially expressed genes by cluster from 5’ scRNA-seq.

Table SI3: Related to Figure 6. Network propagation analysis results.

