## Supplemental figures for "Hepatitis C Virus Infects and Perturbs Liver Stem Cells"

**This PDF file includes:**

Figures S1 to S6
Table S1
Captions for Supplementary Items SI1 to SI3

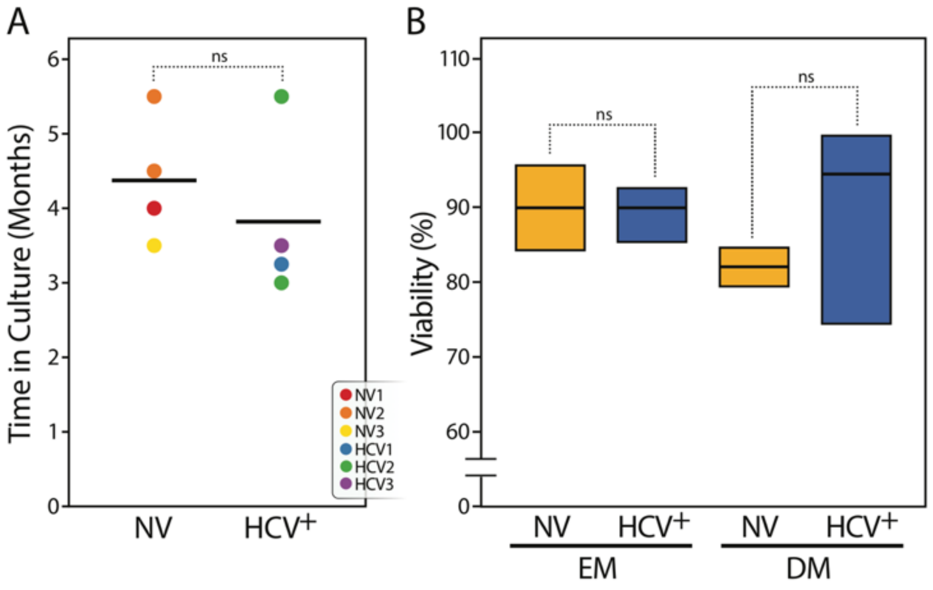


**Figure S1. Related to Figure 1. EM Organoids have Similar Lifespans and Viability between HCV Positive and Negative Donors. (A)** Time in culture is plotted for NV and HCV+ organoids. Organoid termination in culture was recorded by no organoid outgrowth followed passaging. Organoids were grown at two separate times from input tissue for HCV2 and NV2, but only once for all other donors. Horizontal lines represent the grand mean. **(B)** Viabilities of EM and DM day 10 organoids measured by trypan blue staining. Organoid viabilities were grouped according to donor infection status and EM or DM and represented as mean ± SEM.

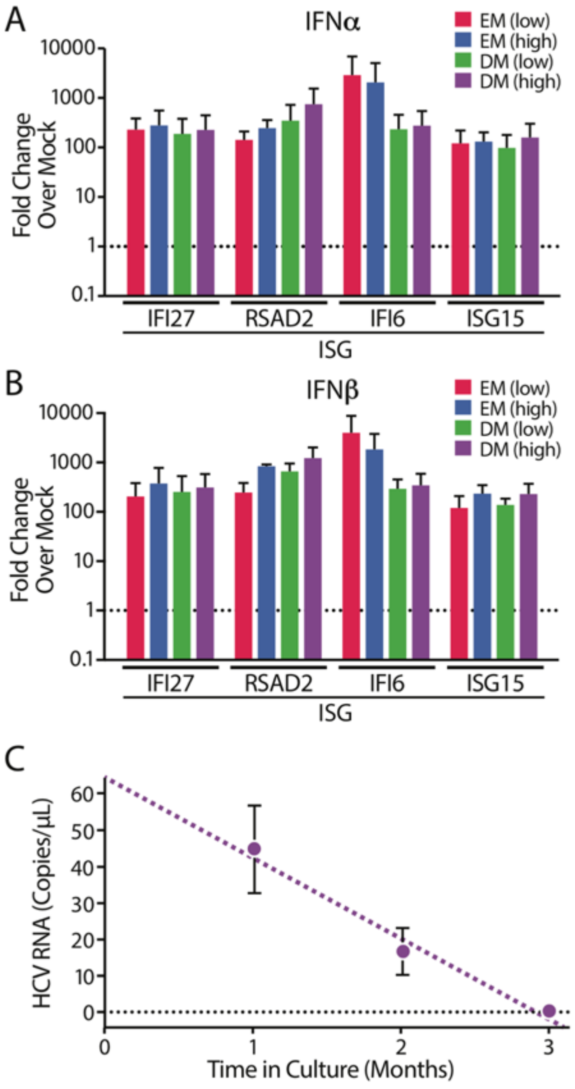


**Figure S2. Related to Figure 2. Liver Organoids are Responsive to Interferon Treatment.** (A) RT-qPCR of interferon stimulated genes (ISGs) treated with low = 500 U/ml or high = 5000 U/ml of (A) IFNα or (B) IFNβ for 18hrs. For each transcript, expression was normalized to 18S and plotted as a fold change over the gene's expression in mock-treated samples ($\Delta\Delta C_T$) which was set to 1 and is represented by a dotted line. Fold change is plotted on a logarithmic scale. Data is from n=2 stimulation experiments and represented as mean ± SD. (C) ddPCR of HCV RNA in HC1 EM organoids over time. Each time point has at least n=3 biological replicates and is represented as mean ± SD.

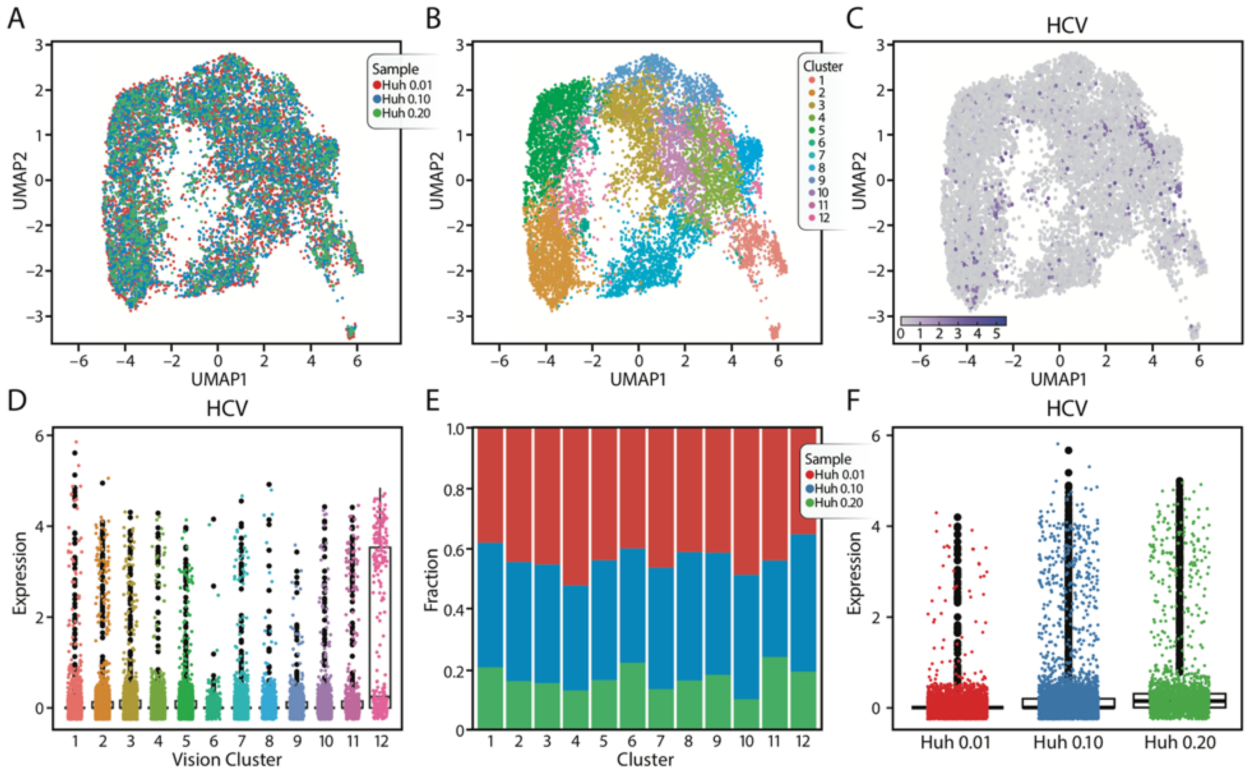


**Figure S3. Related to Figure 3 and Figure 4. HCV Infection of Hepatoma Cells (A)** UMAP plotted by sample of 5' scRNA-seq with HCV capture oligo performed in Huh 7.5 cells infected for 7 days with a JC1 HCV strain at three MOI (0.01, 0.1, 0.2). **(B)** The same UMAP in (A) is plotted by the 12 clusters identified in scVI. **(C)** UMAP shows HCV RNA expression (as pseudolog) is distributed across all clusters. **(D)** A box plot shows HCV RNA expression on the y-axis (as pseudolog) with cluster on the x-axis. **(E)** A stacked barplot shows the composition by sample of each of the 12 clusters. Clusters are shown on the x-axis and the fraction of cells from each donor within that cluster is plotted on the y-axis. **(F)** A box plot shows HCV RNA expression on the y-axis (as pseudolog) with Huh 7.5 sample on the x-axis.

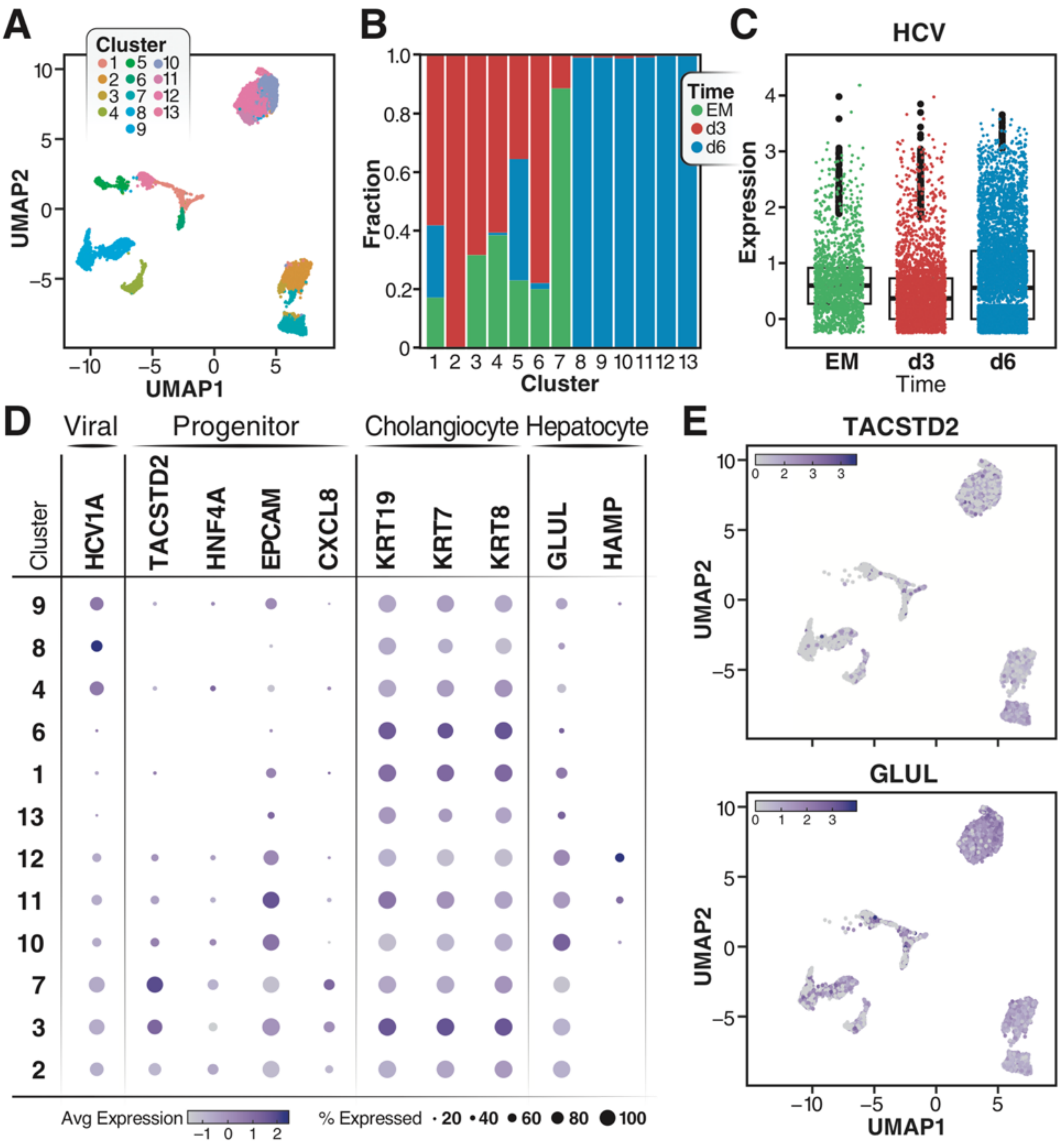


**Figure S4. Related to Figure 4, 5, and 6. Uninfected cells differentiate into both hepatocyte and ductal like cells (A)** A UMAP shows the 13 clusters identified in Figure 4 including the location of cluster 5 **(B)** A stacked bar plot (as pseudolog) shows the composition by sample of each of the 13 clusters identified in scVI. Clusters are shown on the x-axis and the fraction of cells from each donor within that cluster is plotted on the y-axis. **(C)** A box plot shows HCV RNA expression on the y-axis (as pseudolog) across the three HCV1 samples on the x-axis. **(D)** A dot plot shows gene expression (as pseudolog) of progenitor, ductal and hepatocyte markers across the 13 clusters. **(E)** UMAPs show gene expression (as pseudolog) for progenitor marker, TACSTD2, and hepatocyte marker, GLUL.

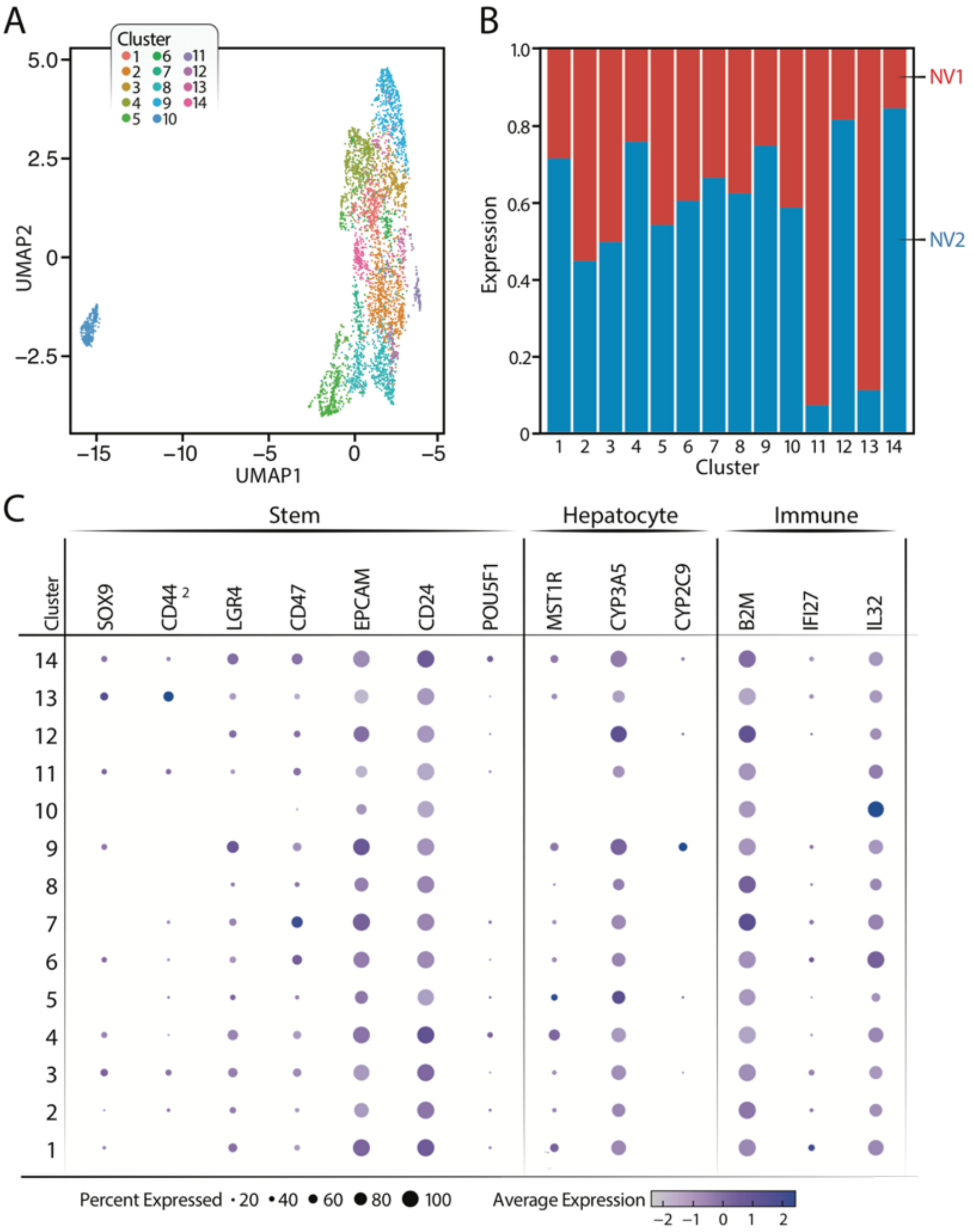


**Figure S5. Related to Figure 4. Differentiated cells from Non viral donors do not have clusters similar to HCV$^{high}$ clusters (A)** UMAPs are generated from 3' scRNA-seq data for DM day 10 organoid samples from donor NV1 and NV2 and plotted according to the 14 clusters identified in scVI. **(B)** A stacked bar plot shows the composition by sample of each of the 14 clusters. Clusters are shown on the x-axis and the fraction of cells from each donor within that cluster is plotted on the y-axis. **(C)** A dot plot shows gene expression (as pseudolog) for stem cell, hepatocyte, and immune markers identified in 5' scRNA-seq data for HCV1 organoids.

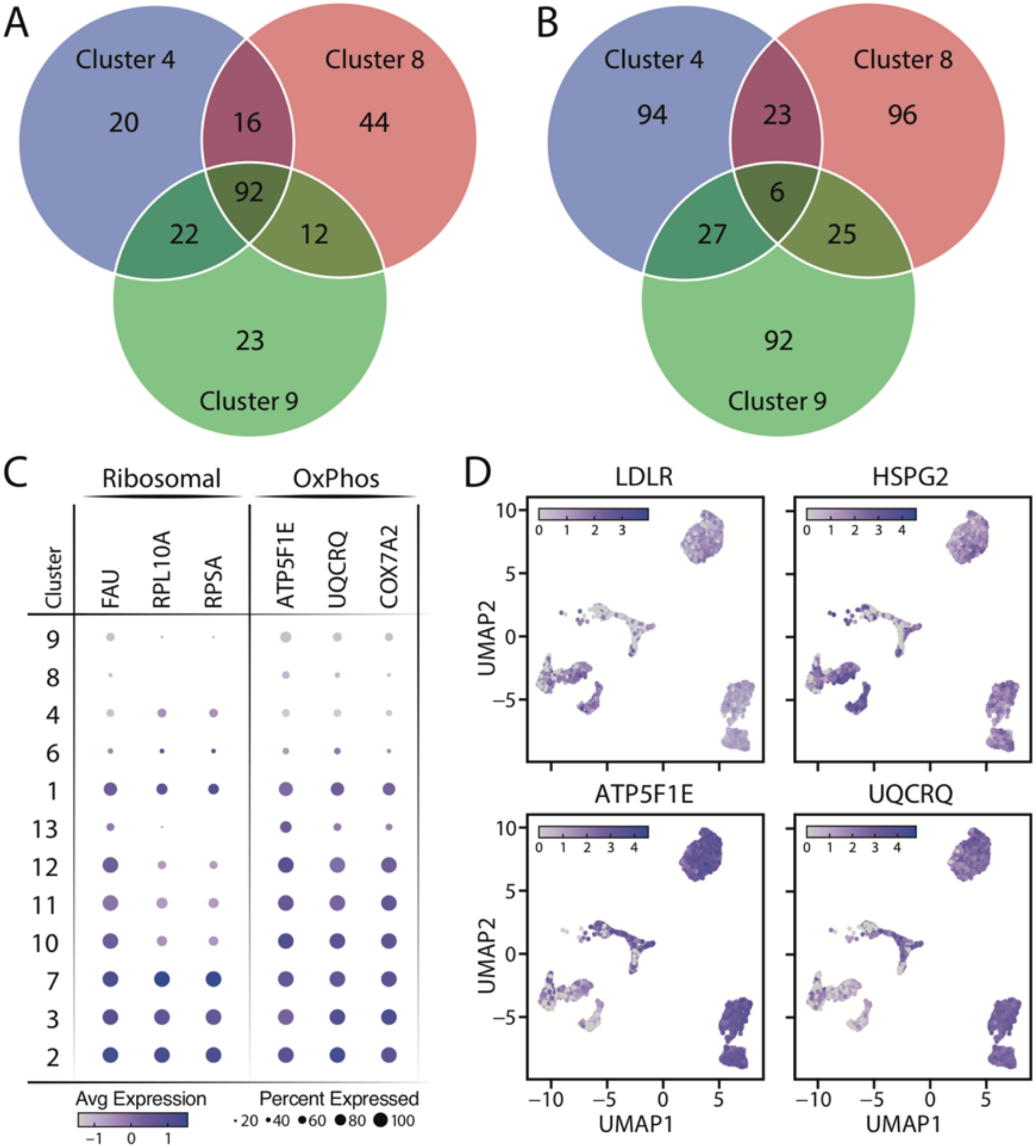


**Figure S6. Related to Figure 5 and 6. HCV Infection of Liver Stem Cells Down-Regulates Oxidative Phosphorylation.** (A) Venn Diagram of the top 150 gene up-regulated genes in HCV$^{high}$ clusters 4, 8 and 9 from Fig. 4 (B) Venn Diagram of the top 150 gene down-regulated genes in HCV $^{high}$ clusters 4, 8 and 9 from Fig. 4. (C, D) A dot plot (C) and UMAP plots (D) are shown for genes in different GO terms, including oxidative phosphorylation (OxPhos) and extracellular matric (matrix). UMAP lots are also shown for hepatocyte/lipid metabolism markers LDLR and HSPG2.

| Sample Name | Library Kit | Cells | Reads per Cell | Genes per Cell | Median UMI per Cell | Sequencer |
|---|---|---|---|---|---|---|
| NV1 EM | 3’ | 2,603 | 25,760 | 2,399 | 8,017 | HiSeq 4000 |
| NV2 EM | 3’ | 1,971 | 95,843 | 4,005 | 20,336 | NovaSeq 6000 |
| HCV1 EM | 3’ | 2,702 | 73,456 | 4,435 | 21,758 | HiSeq 4000 |
| HCV2 EM | 3’ | 2,848 | 43,584 | 3,309 | 13,252 | HiSeq 4000 |
| HCV3 EM | 3’ | 2,835 | 41,098 | 2,823 | 9,654 | HiSeq 4000 |
| NV1 DM d10 | 3’ | 1,795 | 34,531 | 2,459 | 7,675 | HiSeq 4000 |
| NV2 DM d10 | 3’ | 2,746 | 70,761 | 3,008 | 9.847 | NovaSeq 6000 |
| HCV1 EM | 5’ | 1,620 | 292,982 | 4,635 | 24,635 | NovaSeq 6000 |
| HCV1 DM d3 | 5’ | 2,898 | 187,140 | 4,282 | 21,792 | NovaSeq 6000 |
| HCV1 DM d6 | 5’ | 5,567 | 92,391 | 2,282 | 19,981 | NovaSeq 6000 |
| Huh 7.5 MOI = 0.01 | 5’ | 5,352 | 278,035 | 6,159 | 38,210 | NovaSeq 6000 |
| Huh 7.5 MOI = 0.1 | 5’ | 4,732 | 264,293 | 6,286 | 42,120 | NovaSeq 6000 |
| Huh 7.5 MOI = 0.2 | 5’ | 1,874 | 758,599 | 6,912 | 51,727 | NovaSeq 6000 |

**Table S1. Related to Figures 2, 4, and 5. Single-Cell RNA-sequencing Statistics for Liver Organoid Samples.**

**Table SI1: Related to Figure 2. Differentially expressed genes by cluster from 3' scRNA-seq.** All significantly differentially expressed genes in each cluster from 3′ scRNA-seq of EM organoids from five donors in Figure 2.

**Table SI2: Related to Figure 4 and 5. Differentially expressed genes by cluster from 5' scRNA-seq.** All significantly differentially expressed genes in each cluster from HCV1 EM, and day 3 (d3) or day 6 (d6) DMs based on 5′ scRNA-seq with HCV capture oligo in Figure 4.

**Table SI3: Related to Figure 6. Network propagation analysis results.** Detailed results from network analysis of commonly up- or down-regulated genes from $HCV^{high}$ clusters in HCV1 EM, and day 3 (d3) or day 6 (d6) DMs based on 5′ scRNA-seq with HCV capture oligo in Figure 4 and previously published HCV interactome.
